# Capturing changes to animal complexity from quantifiable patterns in genomic data

**DOI:** 10.1101/2024.08.22.609214

**Authors:** Kevin J. Peterson, Alexander W. Clarke, Grygoriy Zolotarov, Bradley Deline, Mark A. McPeek, Pedro Martinez, Bastian Fromm

## Abstract

A prevailing problem in evolutionary biology is elucidating the “genotype-phenotype map” that characterizes how genomic activities regulate different aspects of organismal morphology and their variability in both space and time. Here, we explore potential causality between genome content and both morphological complexity and disparity by compiling the regulatory components (i.e., transcription factors, RNA binding proteins, and microRNA families) as well as a representative set of non-regulatory “housekeeping genes” in 32 species belonging to a wide variety of animal phyla, altogether encapsulating a number of varying genomic characteristics and morphological diversities. A principal component analysis of these four non-overlapping genomic components from each of these 32 species in relation to their last common ancestor revealed that no relationship exists between genome space and disparity, as changes to animal body plans appear to be largely the result of changes to the gene regulatory networks that govern animal development rather than gaining or losing specific sets of regulatory genes. However, using both phylogenetically correlated as well as phylogenetically uncorrelated statistical tests, we find a strong relationship between the loss of all considered gene types and the advent of some parasitic taxa, as well as between microRNA innovations and organismal complexity. While this analysis of genomic features suggests how complexity and disparity are each encoded in the genome, further analysis of the regulatory networks in which they participate should provide a more comprehensive description of how organisms diversify their morphologies over time through alterations in their genomic components.

**SIGNIFICANCE STATEMENT:** Variation in morphological form can be measured by considering either disparity (i.e., the amount of variance in morphological form) or complexity (i.e., the difference in the number of their “parts” including genes or cells). To date, discerning the genomic basis of either has remained largely unknown. Here, we compile the regulatory genome of the last common ancestor of bilaterian metazoans, including its transcription factors, RNA-binding proteins and microRNA families, and ask how changes to these repertoires potentially reflects changes in either disparity or complexity. We find that although there is no relationship between the regulatory genome and disparity, there is a robust relationship between the loss of regulatory genes relative to housekeeping genes, especially in parasitic taxa. Further, there is a strong relationship between increases to microRNAs and complexity, which likely reflects the unique role microRNAs play in increasing both the accuracy and the precision of gene expression during development.

## Introduction

Understanding macroevolutionary trends in animal evolution requires knowing the causal relationship between genomic and morphological variation, otherwise known as the genotype to phenotype map (Lewontin 1974). There are two primary metrics that assess morphological variation at the species level, disparity and complexity (McShea & Brandon 2010). Disparity (or morphological diversity) is a metric that quantifies the variation among taxa in terms of the position and variance within morphospace, as opposed to taxonomic diversity that measures the number of discrete taxonomic units at any rank in the Linnean hierarchy (Runnegar 1987; Foote 1997; Erwin 2007). Complexity, on the other hand, is an assessment of the number of “parts” that make up an organism (McShea 2000; McShea & Brandon 2010), where the number of parts can be whatever metric the investigators choose and at any hierarchical level, whether genes, cells or organismal-level structures.

To date, only a single study has attempted to explore disparity across the entire animal kingdom, and to test for causality by looking for statistical correlates with other facets of variation, including taxonomic diversity, organismal-level complexity and genomic complexity. Deline et al. (Deline et al. 2018) assembled a morphological data set that consisted of 1,767 characters from 212 extant, terminal taxa belonging to 34 animal phyla. They considered a number of potential correlates to the resulting disparity and found that several of these metrics were strongly correlated, including species-level diversity, body size, and the estimated number of cell types in the adult animal (Valentine et al. 1994), the now standard metric for assessing organismal complexity in metazoans (Bell & Mooers 1997; Lee et al. 2007; Jiang & Xu 2010; Vogel & Chothia 2006). Further, they showed a strong correlation between disparity and genome length, as well as to the total number of miRNA families sampled from at least one representative species per analyzed phylum, but not to any aspect of protein-level complexity including the repertoire of protein domains or protein-domain architectures. Because of these results, Deline et al. (Deline et al. 2018) argued that expansions in gene regulatory complexity could potentially underlie the evolution of metazoan morphological complexity, which then positively impacts morphological disparity as an increase in the number of cell types allows for a greater area of morphospace exploration.

However, the Deline et al. (Deline et al. 2018) analysis was necessarily restricted by its taxonomic coarseness (where most taxa were represented at the ordinal level), the interdependence between the reconstruction of character states and the underlying topology of the phylogenetic tree, as well as the limited data for some of the molecular metrics for many of the analyzed taxa. For example, miRNA family-level information was only available for 10 of the 34 animal phyla analyzed, and most of the species (33 of 47) were either chordates or arthropods. Further, aside from gross estimates of the number and type of protein domains, no other aspect of protein-mediated gene regulation was considered. Here, to rectify these limitations, we constructed the “genome-space” for 32 bilaterian species and a “eumetazoan” outgroup (i.e., any cnidarian or placozoan) to capture how the players involved in transcriptional, co-transcriptional, or post-transcriptional gene regulation were gained or lost since these 32 species last shared a common ancestor. These 32 taxa were chosen because of the high quality of their sequenced genomes, the availability of small RNA reads required for the annotation of microRNA repertoires, as well as spanning a range of relevant genomic, morphological and ecological characteristics (Table 1). Using robust statistical methods that control for phylogenetic interdependence (Felsenstein 1985), we show that, although morphological disparity and genome-space are uncorrelated, there is a strong and unexcepted correlation between the loss of regulatory and housekeeping genes that is simply exacerbated in some of the analyzed parasitic species and, as long hypothesized (Sempere et al. 2006), a striking relationship between the number of miRNA families and organismal-level complexity.

**Table 1.**
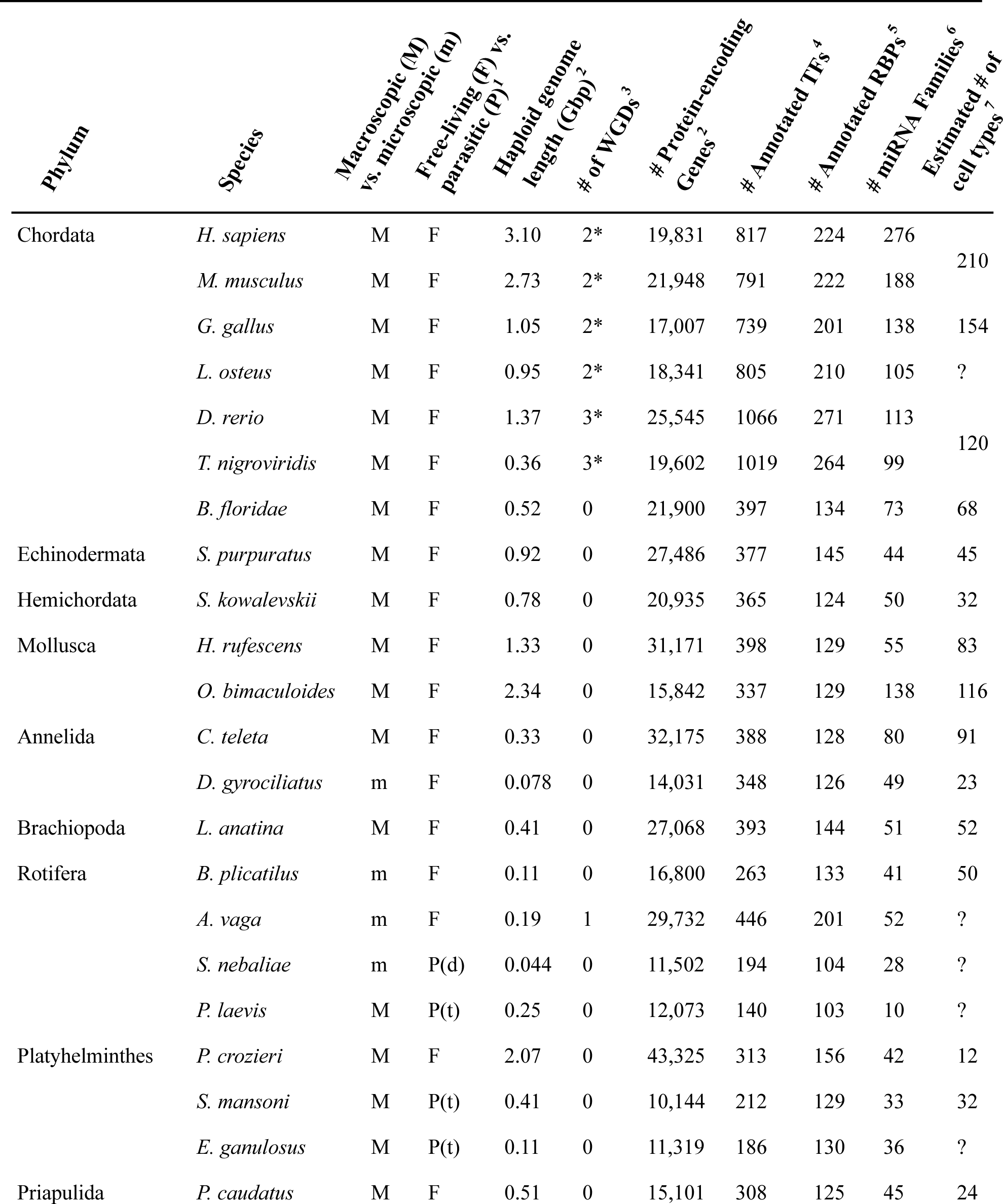
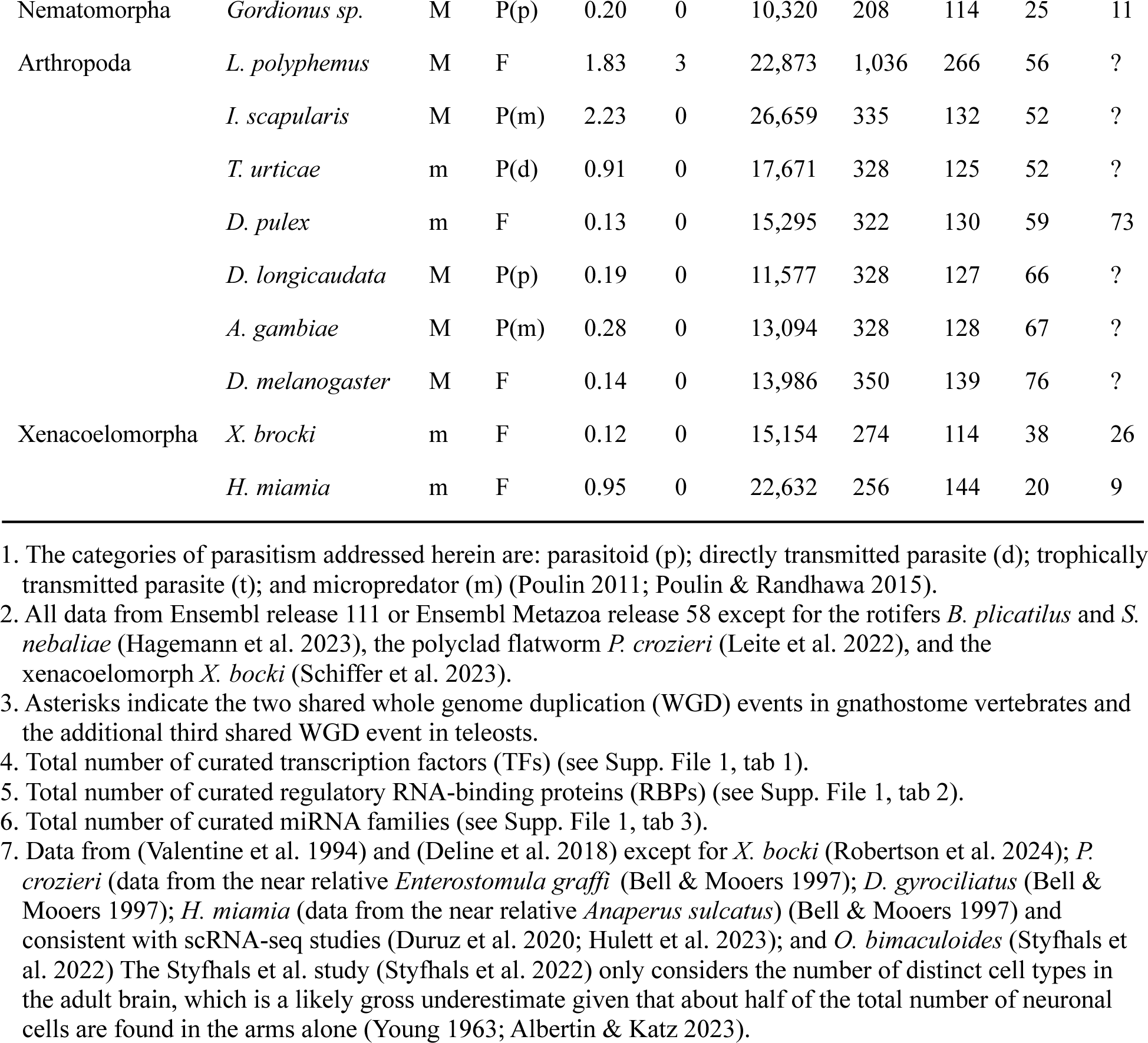
The 32 species and the variables under consideration.

## Results

To explore potential connections between genomic variation and species-level disparity as well as organismal-level complexity, we compiled four non-overlapping genomic data sets from 32 bilaterian species and a “eumetazoan” outgroup. Three of these data sets captured gene regulation at the transcriptional, co-transcriptional, and post-transcriptional steps, plus a fourth data set that considered single-copy housekeeping genes present in the metazoan last common ancestor (the BUSCO gene set, (Manni et al. 2021) (Supp. File 1). We proceeded as follows. First, starting with the list of human transcription factors (TFs) (Messina et al. 2004), each of these genes were searched using the Ensembl “orthologue” function in three other vertebrate species: the mouse *Mus musculus*, the chicken *Gallus gallus* and the spotted gar *Lepisosteus oculatus*. Any gene found in at least three of the four species, plus all of its paralogues as found using the “paralogue” function in Ensembl, where then each mapped to one of the four gnathostome-specific sub-genomes (Lamb 2021; Simakov et al. 2020) (Supp. File 1, tab 1). This allowed us to distinguish between paralogues generated by the two whole-genome duplication (WDG) events in early vertebrate history from potential orthologues shared with one or more invertebrate taxa.

Next, we used a combination of reciprocal blastp (Tatusov et al. 1997) and phylogenetic analysis against the publicly available predicted proteins from the sequenced genomes of a select set of macroscopic invertebrate taxa whose genomes are particularly complete and that did not undergo any WGD events: the cephalochordate *Branchiostoma floridae* (Putnam et al. 2008), the hemichordate *Saccoglossus kowakevskii* (Simakov et al. 2015), the sea urchin *Strongylocentrotus purpuratus* (Consortium et al. 2006), the polychaete annelid *Capitella teleta* (Simakov et al. 2013), the brachiopod *Lingula anatine* (Luo et al. 2015), and the abalone *Haliotis rufescens* (Griffiths et al. 2022). From this, we determined that 330 single-copy transcription factors were present in their last common ancestor (LCA). This estimate ignores TF families where orthology was difficult to infer (e.g., GATA, DMRT, THAP and virtually all C2H2 zinc-finger genes minus a select few that encode products important in neural development and with discernable orthology amongst the considered taxa), as well as genes belonging to well-characterized families, but where the single-copy status in this LCA was difficult to establish or where orthology was difficult to discern (see Supp. file 1, tab 1).

This manually-curated list of 330 TFs present in this LCA must be an underestimate though because of the known losses of TFs in the human lineage (Takatori et al. 2008). To discover this suite of genes lost in the human lineage, we next characterized the TF repertoires of the fruit fly *D. melanogaster* and the octopus *O. bimaculoides* (see Methods). Again, each protein was reciprocally blasted and phylogenetically analyzed against the genomes of this same suite of invertebrate species. We found an additional 26 TFs that were present in the fly or octopus and at least in one of the three invertebrate deuterostome species analyzed, raising the number of TFs present in their LCA to 356. However, with the exception of zinc-finger containing proteins, because all potential blastp hits in all subject taxa were also explored across this same set of species, an additional five TFs were present in this LCA that were lost in all three query taxa (human, fly, and octopus). Therefore, we curated a total of 361 TFs that were present in the genome of their LCA, with 34 losses and an additional 490 TFs that evolved in the human lineage after it split from the LCA of fruit fly and octopus. Most of these gains (405) are the result of the two rounds of WGD early in vertebrate history (Dehal & Boore 2005; Marlétaz et al. 2024; Yu et al. 2024; Simakov et al. 2020; Lamb 2021). This complement constitutes nearly half of the known or suspected TFs in the human genome, with most of the remaining TFs belonging to the largely eutherian-enriched C_2_H_2_ zinc-finger family (Lambert et al. 2018; Tupler et al. 2001).

Single representative sequences of each of these 361 TFs, in addition to any TF present in one or more invertebrate taxa that evolved after their LCA, were assembled into a fasta file (Supp. File 2) and used to query 20 additional bilaterian species whose biology allowed us to explore a range of adult sizes, ecologies, and complexities (Table 1). Size, increases to which have long been suspected as an important contributor to increases in complexity as well as disparity (Bonner 2004; Bell & Mooers 1997; Carroll 2001; Deline et al. 2018), was divided into just two categories, macroscopic (>∼1mm in adult length) versus microscopic. The microscopic category included a likely example of miniaturization (the crustacean *Daphnia pulex*, (Ye et al. 2017), progenesis (the polychaete annelid *Dimorphilus gyrociliatus*, (Martín-Durán et al. 2021), neoteny (the rotifers *Brachionus plicatilus*, *Adineta vaga*, and *Seison nebaliae*, (Hagemann et al. 2023), and two instances of potentially primitively small size, the xenacoelomorphs *Xenoturbella bocki* (Schiffer et al. 2023) and *Hofstenia mimia* (Gehrke et al. 2019).

Differences in morphological complexity were addressed by examining species covering a range of cell types in the adult, the accepted metric for complexity assessments in metazoans (Deline et al. 2018). Vertebrates and the octopus (Styfhals et al. 2022) have well over a hundred different cell types in their brains alone, whereas animals like the free-living polyclad flatworms including *Prostheceraeus crozieri* (Leite et al. 2022) have only 12 described cell types in the entire adult worm (Table 1). Differences in genomic complexity were addressed by comparing species spanning both a range of genome sizes as well as gene complements. Indeed, because whole genome duplications have long been invoked to explain morphological complexity (Ohno 1970), we examined four lineage-specific sets of whole genome duplication (WGD) events: the two rounds of WGD in vertebrates that occurred after the split from the cephalochordate *B. floridae*, but before the split between *L. oculatus* and *H. sapiens* (Dehal & Boore 2005; Simakov et al. 2020; Lamb 2021; Yu et al. 2024; Marlétaz et al. 2024); the third round of WGD that occurred in the teleost lineage before the last common ancestor of the zebrafish *Danio rerio* and the pufferfish *Tetraodon nigroviridis* but after it split from the basal actinopterygians including the spotted gar *L. oculatus* (Braasch et al. 2016; Amores 1998; Jaillon et al. 2004); the single round of WGD that characterizes bdelloid rotifers including *Adineta vaga* (Welch et al. 2008; Simion et al. 2021); and lastly the three rounds of WGD that characterizes living horseshoe crabs including *Limulus polyphemus* (Kenny et al. 2016; Nong et al. 2021).

Finally, because parasites are often thought of as simplified – both morphologically as well as genomically – relative to their free-living counterparts, we examined the regulatory repertoires of nine different parasitic taxa belonging to four of the six established evolutionary strategies of parasitic species as proposed by (Poulin 2011; Poulin & Randhawa 2015). Parasitoids, which grow inside a single host and kill that host as a normal and necessary part of their development, included the wasp *Diachasmimorpha longicaudata* and the nematomorph *Gordionus sp.* The directly-transmitted parasites, which infect only one host individual during their lifetime, included the rotifer *Seison nebaliae* (Mauer et al. 2021) and the spider mite *Tetranychus urticae* (Grbić et al. 2011). The micropredators, which on the other hand feed on multiple host individuals during their lifetimes, included the mosquito *Anopheles gambaie* (Neafsey et al. 2015; Holt et al. 2002) and the tick *Ixodes scapularis* (De et al. 2023). Finally, we profiled three trophically-transmitted parasites, the tape worm *Echinococcus granulosus* (Zheng et al. 2013; Tsai et al. 2013), the liver fluke *Schistosoma mansoni* (Berriman et al. 2009), as well as the acanthocephalan *Pomphorhynchus laevis* (Hagemann et al. 2023; Mauer et al. 2020), all of which negatively affect the fitness of at least two different host individuals belonging to two different animal species. Each of these instances of parasitism are known (Weinstein & Kuris 2016) or suspected (Brabec et al. 2023) to have evolved independently of one another within each of the four considered parasitic strategies.

To capture potential changes to the machinery that controls co-transcriptional gene regulation, a second manually-curated dataset of RNA-binding proteins (RBPs) was assembled (Supp. File 3). Starting with the census of RBPs in the genomes of the human (Gerstberger et al. 2014) and the fruit fly (Gamberi et al. 2006), and using a similar approach to the assembly of TFs described above, we curated 124 RBPs in each of the 32 species that were present in the their LCA with GO terms in either human or fruit fly that indicate involvement in the regulation of (alternative) splicing, the regulation of the export or stability of mature mRNA, or the (positive) regulation of translation (Supp. File 1, tab 2). As expected (Kerner et al. 2011), virtually all of these regulatory RBPs (123 of 124) were inherited from the eumetazoan LCA with very few novel regulatory RBP genes evolving after this LCA in any analyzed species. Indeed, we found only 6 regulatory RBPs evolving in the human lineage including one shared with the cephalochordate (*Zc3h7*), and two novel RBPs in the fruit fly lineage with one (*sex lethal*) primitive for protostomes and the second (*mextli*) restricted to the analyzed ecdysozoans. Moreover, *H. sapiens* retains a complete repertoire of regulatory RBPs with no losses from what it inherited from this LCA.

The negative regulation of translation was analyzed by assessing the gains and losses of 763 miRNA families found in at least one of the 32 queried species as curated in MirGeneDB (Fromm et al. in prep.) (Supp. File 1, tab 3). As is well known (Tarver et al. 2018, 2013; Fromm et al. 2022; Fromm 2024), and unlike both transcription factors and especially RBPs, most miRNA families evolved after their LCA as only 32 of the 763 miRNA families were present in this ancestral population. RBPs known (e.g., LIN28, TRIM71) or suspected (e.g., PRKRA/TARBP2) to regulate miRNA function (Treiber et al. 2017) were also considered in our tally of regulatory RBPs (Supp. File 1, tab 2). Altogether, over 21,000 regulatory genes or gene families were curated for these 32 taxa. Finally, to gauge how any changes to the regulatory portion of the genome compare to its non-regulatory component (i.e., the “housekeeping” repertoire of genes), we used the curated 954 BUSCO gene data set (Manni et al. 2021; Simão et al. 2015) for each of the 32 sampled species (Supp. File 1, tab 4). Eleven of these BUSCO genes overlapped either the TF file or the RBP file and thus were removed from the BUSCO file generating a dataset of 943 non-regulatory and non-redundant housekeeping genes (Supp. File 1, tab 5).

To construct an image of genome-space for these 32 species, we characterized the percentage gains and losses of all considered TFs, RBPs, miRNA families, and BUSCO genes relative to the LCA. But, because of the controversy surrounding the phylogenetic position of the xenacoelomorphs – whether they are basal bilaterians or instead allied with the ambulacrarian deuterostomes (i.e., echinoderms and hemichordates) – we constructed two different ancestral repertoires. The first dataset considered them to be a monophyletic group of basal bilaterians (Cannon et al. 2016; Juravel et al. 2023; Rouse et al. 2016) and as such any gene present in either any non-bilaterian, *X. bocki* or the acoel *H. miamia* was tallied as present in the bilaterian LCA (hereafter referred to as the LCB or last common bilaterian ancestor) (Supp. File 1, tabs 6, 7). The second dataset considered the xenaceolomorphs deuterostomes allied with the ambulacrarians (Philippe et al. 2011; Mulhair et al. 2022), and thus any gene present in at least one protostome species and one deuterostome species was reconstructed as present in their LCA, known as the last common nephrozoan ancestor or LCN (Supp. File 1, tabs 12, 13). Because the results were statistically indistinguishable from one another irrespective of their phylogenetic position (Supp. Fig. 1), we only show the results with Xenacoelomorpha reconstructed as the sister taxon to the remaining bilaterians as this necessitates fewer gene losses in the xenacoelomorphs.

To explore any trends in the gains or losses of regulatory genes relative to the LCB, we performed a principal components analysis (PCA) to construct a smaller set of independent variables that captures the main axes of variation in the full dataset (Fig. 1). Nearly 100% of the variation within these four gene categories (TF, RBP, miRNA and BUSCO) can be described just with the first three principal components. Principal components axis 1 (GPC1) explains 92.9% of variation in the full dataset and quantifies primarily gains of miRNA families (largest positive loading) with relatively small contributions from gains of TFs and RBPs (Table 2). The limit of GPC1 relative to the bilaterian LCB (Fig 1, cross) is defined by the position of the six vertebrate species and *O. bimaculoides* due largely to the unique amount of miRNA innovation in each of these lineages not seen elsewhere among bilaterians (Heimberg et al. 2008; Zolotarov et al. 2022). In fact, within the vertebrates, both mouse and human are extended along GPC1 relative to the three fish species and the chicken due to the dramatic pulse of miRNA innovation that characterizes eutherian mammals relative to all other vertebrates (Tarver et al. 2018; Devor & Peek 2008). In addition, human is extended along GPC1 relative to mouse due to a second pulse of miRNA innovation that occurred within the primates relative to all other eutherian lineages (Langschied et al. 2023).

**Figure 1.**
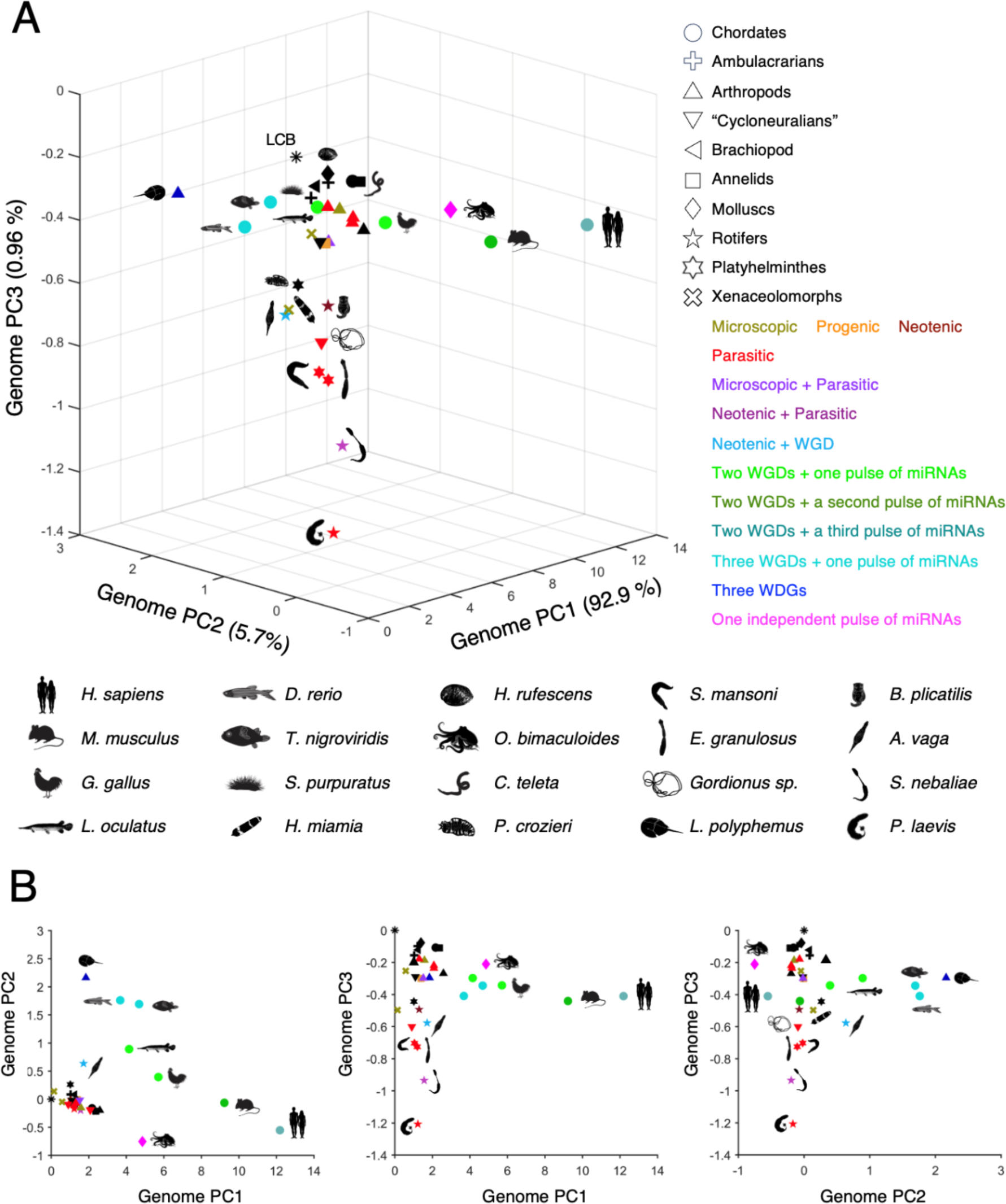
Construction of genome space for 32 bilaterian species using principal component analysis in relation to the bilaterian last common ancestor (LCB, asterisk). **A.** Three-dimensional view. **B.** The three 2-dimensional views. Notice that that amniotes and the octopus define the limit of GPC1 with respect to the LCB, the three species that underwent three rounds of whole genome duplication (the horseshoe crab *L. polyphemus* and the two teleost fish *D. rerio* and *T. nigroviridis*) define the limit of GPC2, and the four parasites (the acanthocephalan *P. laevis*, the rotifer *S. nebalie*, the trematode *S. mansoni* and the cestode *E. granulosus*) define the limit of GPC3.

**Table 2.** Factor loadings for the first three principal components extracted from the dataset of gains and losses of the BUSCO, TXF, RBPs and miRNA families.

|  |  | **Loadings** |  |  |
| --- | --- | --- | --- | --- |
|  |  | **GPC1** | **GPC2** | **GPC3** |
| **Gains** | BUSCO | 0.0032 | 0.0139 | 0.037 |
|  | TXFs | 0.1657 | 0.8664 | 0.0222 |
|  | RBPs | 0.0787 | 0.4582 | -0.1904 |
|  | miRNA Families | 0.9826 | -0.185 | -0.0141 |
| **Losses** | BUSCO | 0.0141 | 0.0274 | 0.4144 |
|  | TXF | 0.0241 | 0.0539 | 0.5947 |
|  | RBPs | 0.0117 | 0.0257 | 0.2641 |
|  | miRNA Families | 0.003 | 0.0242 | 0.6054 |
| % Explained |  | 92.87 | 5.69 | 0.96 |
| eigenvalues |  | 6.44 | 0.39 | 0.066 |

Principal components axis 2 (GPC2) explains 5.7% of the variation in the original dataset and quantifies gains of both TFs and RBPs (Table 2). The limit of GPC2 relative to the bilaterian LCB is defined by the position of the horseshoe crab *L. polyphemus* and the two teleost species *D. rerio* and *T. nigroviridis*, all of which underwent three rounds of WGD (Amores 1998; Jaillon et al. 2004; Kenny et al. 2016; Nong et al. 2021). Also extended along GPC2, and away from the LCB, is the rotifer *A. vaga* that experienced a single WGD (Welch et al. 2008; Simion et al. 2021), and the two other vertebrate species – *L. oculatus* and *G. gallus* – that share two of the three WGDs found in teleosts (Lamb 2021; Simakov et al. 2020). Of course, these two WGDs are also shared with the eutherian mammals, but the dramatic pulse of miRNA innovation that characterize eutherians relative to all other vertebrates, and simian primates relative to all other eutherians, drive mouse and human away from these taxa down GPC1 (Fig. 1B).

Principal components axis 3 (GPC3) explains just under 1% of the variation in the data set and quantifies the loss of genes belonging to all four gene categories (Table 2). Although all taxa have lost some portion of the ancestral repertoire of TFs and RBPs (Fig. 1), some, like the abalone *H. rufescens*, have lost a relatively small proportion of their ancestral regulatory repertoire. In contrast, the free-living rotifers and the polyclad flatworm have lost a substantial portion of their regulatory repertoire, a trend that continues in their parasitic relatives that, along with the nematomorph, make them outliers on GPC3 (phylogenetic generalized least squares analysis: *F*_1,30_ = 9.08, *P*<0.01). Indeed, we find a strong phenotypic correlation between the number of protein-encoding genes (PEG) and GPC3 (*r*_31_ = 0.37, *P*<0.05), as expected given that parasites, because they can rely on host gene products for many aspects of their biology (Wolf & Koonin 2013; Poulin & Randhawa 2015), are often characterized by gene loss (Jackson 2015; Cunha et al. 2023; Tsai et al. 2013; Coghlan et al. 2019; Herlyn et al. 2024; Albalat & Cañestro 2016). However, one biological process that parasites cannot exploit from their host is developmental gene regulation. Nonetheless, the acanthocephalan *P. laevis*, for example, has lost 68% of its TF repertoire, 31% of its regulatory RBP repertoire and a striking 81% of its miRNA family-level repertoire (Fig. 2A) (Supp. File 1, tab 6). How these animals regulate development despite such a dramatic loss of developmental regulators remains an outstanding question for future investigation.

**Figure 2.**
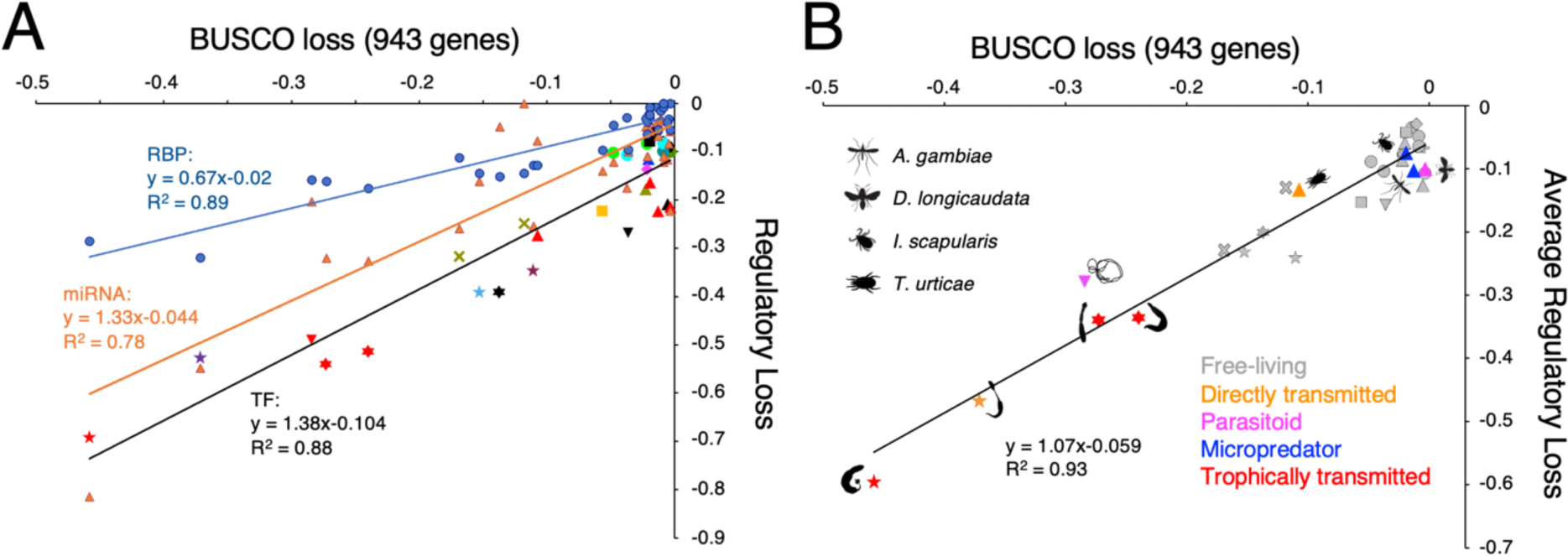
Regression analysis of the loss of the regulatory genes in relation to the loss of BUSCO genes for the 32 sampled species. **A.** The rate of loss of TFs relative to BUSCO gene loss (taxon-specific symbols from Fig. 1, and the black trendline) is statistically indistinguishable from the rate of miRNA family loss (orange) relative to BUSCO gene loss (tests for difference in slope of the regressions: *F*_1,60_ = 0.71, *P*>0.40). RBPs (blue), on the other hand, are lost at approximately half the rate across these taxa in relation to BUSCO relative to the TFs and miRNA families (differences in slope of RBP loss versus the average loss rates of TF and miRNA: *F*_1, 90_ = 15.08, *P*<0.001). **B.** The rate of regulatory loss (the mean percentage of all three gene types) is strongly correlated to the loss of BUSCO genes in both parasitic (colored symbols) as well as free-living (gray symbols) species. Further, the pattern of gene loss does not characterize any particular parasitic strategy. Instead, the only apparent division is between the arthropod versus the non-arthropod parasitic species with the former nested within the analyzed free-living species.

In contrast to the nematomorph and the flatworm and rotiferan parasites, the arthropod parasites are nested within the cloud of points defined by species with a free-living ecology, including taxa sharing the same parasitic strategy with these greatly reduced taxa. For example, despite both taxa being parasitoids, the average regulatory loss for the nematomorph *Gordionus sp.* is 28%, whereas the wasp. *D. longicaudata* has only lost 10% of its ancestral regulatory repertoire (Fig. 2B, magenta), the same as the teleost fish *T. nigroviridis* and a slightly higher retention rate than found in the fruit fly *D. melanogaster* (Supp. Table 1, tab 8). Of the arthropod parasites analyzed, only the mite *T. urticae* shows a moderate reduction in regulatory gene content (13.3%), but this is still a much greater retention than the second directly-transmitted parasite considered, the rotifer *S. nebaliae*, which lost 47% of its regulatory genes (Fig. 2B, orange).

A surprising result is that for both parasitic and free-living taxa, the trend for gene loss across the 32 analyzed species is the same (test for difference in slope between parasites versus non-parasites is F_1,28_ =0.01, P>0.90, and test for difference in intercept between the two groups is F_1,29_ = 0.01, P>0.92) (Fig. 2B). Thus, any trend to lose genes in a parasitic species follows the same rules as their free-living counterparts. Indeed, we find that the rate of TF loss and the rate of miRNA family-level loss is statistically indistinguishable from one another when compared to the rate of BUSCO loss (Fig. 2A) (tests for difference in slope of the regression: *F*_1,60_ = 0.71, *P*>0.4). In contrast, the regulatory RBPs were lost at approximately half the rate as compared to the TFs and miRNA families (Fig. 2A, differences in slope of RBP loss versus the average loss rates of TF and miRNA: *F*_2,90_ = 15.08, *P*<0.001). This striking relationship of gene loss between miRNAs and TFs might be explained simply by the fact that the primary targets of miRNAs are, in fact, TFs (Stark et al. 2005; Zare et al. 2014). Why genes that encode RBPs are lost at a slower rate than either transcriptional or post-transcriptional processes remains an open question, but we note that the “housekeeping” RBPs present in the BUSCO gene set are lost at the same rate as regulatory RBPs (test for differences in slope of RBP classes regressed on BUSCO gene set: *F*_1,30_ = 1.62, *P*>0.20). This strongly suggests that there are tighter constraints to losing RBPs – whether non-regulatory or regulatory - relative to TFs and miRNA families, highlighting the central role that RNA metabolism plays in the cell irrespective of whether the species itself is free living or parasitic (Anantharaman et al. 2002).

To understand any potential causality between genome content and morphological variation – whether complexity or disparity – strong correlative support between the two must first be found (McShea 1996). To that end, we repeated the disparity analysis of Deline et al. (Deline et al. 2018) for the 32 species considered herein (Fig. 3) (Supp. File 1, tab 9; Supp. File 4). Our results are largely concordant with Deline et al. (Deline et al. 2018), with the seven arthropod species and the six vertebrate species all defining the limits of two of the three morphological principal coordinate axes; the third is defined by the position of the octopus and the sea urchin (Fig. 3B). Deline et al. (Deline et al. 2018) also found a robust correlation between disparity and the number of miRNA families. Our results at the phenotypic level (i.e., without phylogenetic correction) confirm their findings, with GPC1 correlated with both MPC1 (*r*_31_ = - 0.47, *P*<0.01) and MPC2 (*r*_31_ = −0.61, *P*<0.0005) (Table 3). These authors also found a strong relationship between miRNA families and complexity as measured through the estimated number of cell types (CT), a result we robustly confirm (*r*_20_ = 0.94, *P*<0.001) (Fig. 4A). Finally, we also found a correlation between genome length and disparity as genome length (Gbp) is positively correlated with MPC3 (*r*_31_ = 0.41, *P*<0.05), as well as with GPC1 (*r*_31_ = 0.66, *P*<0.0001). However, complexity is uncorrelated with GPC2, which is dominated by the gains of both TFs and RBPs resulting from whole genome duplication events (*r*_20_ = 0.08, *P*>0.70) (Fig. 4B), and is also uncorrelated with organismal size (all phylogenetical generalized least squares comparisons of macroscopic and microscopic taxa with *P*>0.75).

**Figure 3.**
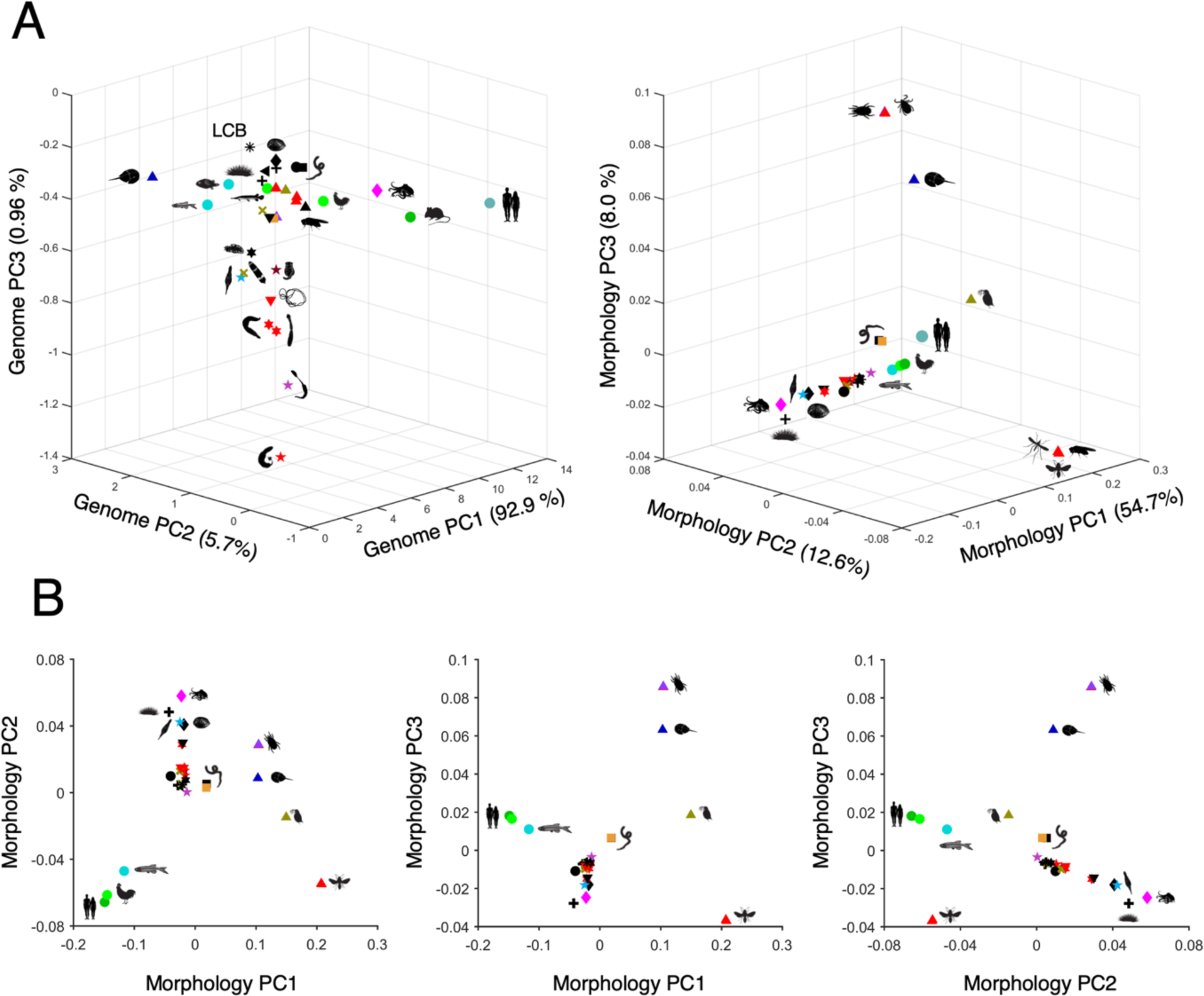
Construction of morphospace for 32 bilaterian species using principal coordinate analysis. **A.** Three-dimensional view in comparison to the genome space from Figure 1A. **B.** The three 2-dimensional views. Here, the arthropods define the limit of MPC1, the vertebrates MPC2, and the octopus and sea urchin (*S. purpuratus*) MPC3. The LCB is clustered within the cloud of data points in the middle (i.e., the “0” value) of each of the axes.

**Figure 4.**
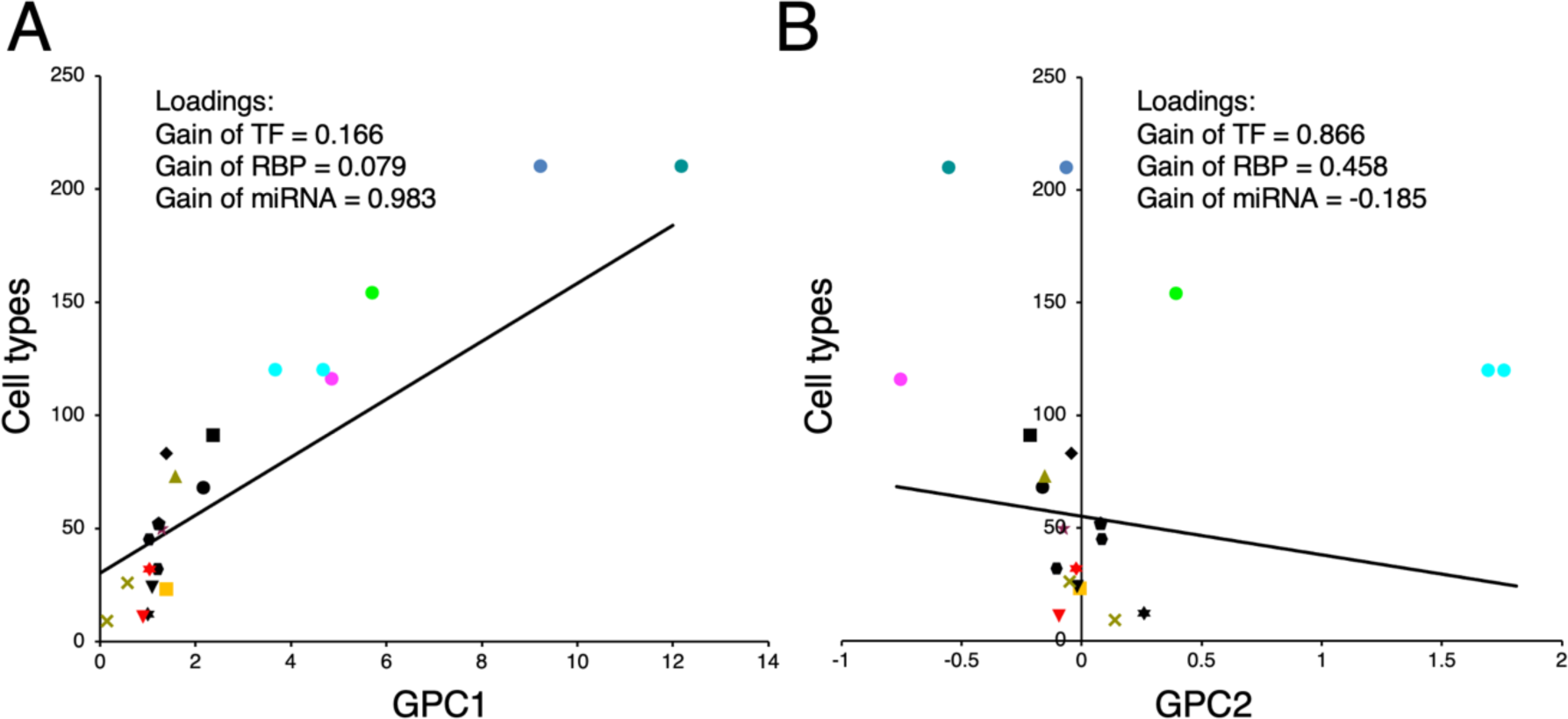
Complexity is strongly correlated with GPC1, which is primarily loaded by gains of microRNA families (**A**, *r*_20_ = 0.94, *P*<0.0001), and is not correlated at all with GPC2, which is primarily loaded by gains of TFs and RBPs generated during WGD events (**B**, *r*_20_ = 0.081, *P*>0.70) (see also Tables 2 and 3). The lines fitted through the data in each panel is the phylogenetic regression of the number of cell types on each of the two genomic PCs. For Panel A, the phylogenetic regression line is Cell Type = 30.40+12.79*GenomePC1 (test for slope=0, *t*_19_=5.04, *P*<0.0001). For panel B, the line is Cell Type = 55.15-17.07*GenomePC2 (test for slope=0, *t*_19_=-1.01, *P*>0.30).

**Table 3.** Correlations between the genome principal components axes (GPC, Fig.1), morphology principal components axes (MPC, Fig. 3), and the biological metrics addressed herein^1^.

|  | GPC1 | GPC2 | GPC3 | MPC1 | MPC2 | MPC3 | Gbp | PEG | CT |
| --- | --- | --- | --- | --- | --- | --- | --- | --- | --- |
| GPC1 | - | -0.007 | 0.045 | -0.473<sup>2</sup> | -0.608 | 0.149 | 0.660 | 0.009 | 0.938 |
| GPC2 | -0.388 | - | 0.010 | -0.197 | -0.203 | 0.330 | 0.047 | 0.244 | -0.081 |
| GPC3 | -0.122 | -0.013 | - | 0.206 | -0.036 | 0.079 | 0.194 | 0.371 | 0.028 |
| MPC1 | 0.002 | -0.079 | 0.423 | - | 0.059 | -0.022 | -0.332 | -0.204 | -0.552 |
| MPC2 | -0.314 | -0.079 | 0.113 | -0.214 | - | 0.015 | -0.120 | 0.190 | -0.317 |
| MPC3 | -0.016 | 0.193 | -0.014 | 0.030 | 0.136 | - | 0.408 | 0.160 | 0.157 |
| Gbp | 0.544 | -0.130 | 0.059 | -0.025 | 0.118 | 0.253 | - | 0.429 | 0.332 |
| PEG | -0.135 | 0.263 | 0.405 | -0.019 | 0.194 | 0.126 | 0.402 | - | 0.665 |
| CT<sup>3</sup> | 0.663 | -0.166 | 0.074 | -0.073 | -0.315 | 0.347 | 0.445 | 0.003 | - |

A potential complicating factor with these results is that the vertebrates are outliers with respect to many of the biological metrics considered herein, including disparity, WGDs (and hence the number of both TFs and regulatory RBPs), miRNA innovation, genome length and the estimated number of cell types (Table 3). Therefore, correlative relationships amongst any of these factors might simply be driven largely by the vertebrates and are not reflective of general trends in animal evolution. To that end, in order to test the robustness of these results, we used phylogenetic independent contrasts (PIC), a statistical approach that circumvents the non-independence of species traits because of their shared evolutionary history, thus removing any correlations between characters due to common ancestry (Felsenstein 1985). The construction of PICs requires the quantitative characters in question, a phylogeny of the species under investigation, and, from this topology, estimates of the branch lengths. To that end, we constructed a fifth non-overlapping set of 19 protein sequences derived from single-copy housekeeping genes in the LCB with an aligned length of nearly 11,500 amino acids (Supp. File 1, tab 10; Supp. File 5).

The correlations among the resulting PICs for each variable are shown below the gray diagonal in Table 3. The two correlative relationships found above and by Deline et al. (Deline et al. 2018) between GPC1 (i.e., miRNAs) and disparity, as well as between cell types (i.e., complexity) and disparity, are no longer supported when the phylogeny of the analyzed taxa is taken into consideration (Table 3, yellow cells above the diagonal). However, our PIC analysis still identifies four significant relationships: 1) between GPC1 and genome length (*r*_30_ = 0.54, *P*<0.002); 2) between the number of PEGs and GPC3 (*r*_30_ = 0.41, *P*<0.05); 3) between the number of PEGs and genome length (*r*_30_ = 0.40, *P*<0.05); and 4) most significantly, between GPC1 and the estimated number of cell types (*r*_19_ = 0.66, *P*<0.002) (Table 3, green cells). As with the uncorrected analysis, the number of cell types was not phylogenetically correlated with GPC2 (*r*_19_ = −0.17, *P*>0.45).

## Discussion

To our knowledge, this is the first attempt to map species-level morphological attributes including complexity and disparity to the underlying regulatory apparatus of their genomes, as measured by the gains and losses of regulatory genes in comparison to the LCB. Our statistical analyses assessing potential relationships between morphological variation and genome composition reveals four clear conclusions. First, no general correlative relationship exists between the number of regulatory genes and the disparity of the animal species examined herein. Although Deline et al. (Deline et al. 2018) found a strong relationship between the number of miRNA families and disparity, these results appear to be driven, at least in part, by the phylogenetic overprint of the analysis. When controlling for phylogeny using PIC (Felsenstein 1985), this purported relationship disappears, and thus, disparity is achieved not by the addition (or loss) of regulatory genes, but instead through their interactions that drive development and ultimately morphological diversity. This is consistent with the fact that like we show here (Supp. File 1), most protein-encoding regulatory genes are quite ancient, having evolved long before the diversification of the bilaterian phyla (Fernández & Gabaldon 2020; Kerner et al. 2011; Degnan et al. 2009; Richter et al. 2018; Srivastava et al. 2010; Paps & Holland 2018). Furthermore, numerous studies have shown that morphological differences between species are often the result of subtle changes to the gene regulatory networks that govern development by modulating either the *cis*-regulatory control of the genes that build the morphology of interest and/or their transcriptional effectors (see e.g., (Wray 2007; Carroll 2008; Peter & Davidson 2015; Hill et al. 2021) for reviews), not the evolution of a new regulatory gene in the species of interest.

Second, the loss of regulatory gene in comparison to the BUSCO gene set follows a clear correlative relationship, irrespective of the ecology of the species under consideration, whether free-living or parasitic (Fig. 2). Although previous studies have documented gene loss across the animal tree (Guijarro-Clarke et al. 2020; Fernández & Gabaldon 2020), of the species analyzed herein, only the nematomorphs, rotifers, and the platyhelminths show extensive loss of regulatory genes, which, in the latter two instances, are exacerbated in their parasitic forms. Nonetheless, the *relationship* of this loss to the BUSCO data set is simply an extension to what is also found in free-living forms (Fig. 2). This pattern suggests that at least from a regulatory perspective – either in terms of unique gains or shared losses – nothing unites parasites as a distinct set of species separate from free-living metazoans. Instead, our results suggest that there are clear constraints in terms of regulatory gene loss in relation to the loss of housekeeping genes, whether the taxon is parasitic or not, with some parasites simply extending this conserved trend in relation to their free-living relatives. Indeed, this pattern of gene loss would appear to refute the hypothesis of Albalat and Canestro (Albalat & Cañestro 2016) who described the nature of gene loss as non-stochastic given that it seemed to be governed by either the gene’s function or its genomic location. We find neither to be true with the genes examined herein across this select panel of metazoan species given that we look genome-wide at several different sets of regulatory and non-regulatory genes. Nonetheless, because the handful of examples examined by us pale in comparison to the hundreds of known occurrences of parasitism evolving independently from free-living ancestors in multiple different animal phyla (Weinstein & Kuris 2016), testing the generality and robustness of this pattern remains an outstanding avenue for future research. Nonetheless, what we found would seem to support Wolf and Koonin’s (2013) (Wolf & Koonin 2013) model of biphasic genomic evolution with pulses of regulatory innovation followed by a “clock-like” pattern of loss that is simply exacerbated in some – but certainly not all – parasitic lineages (see also (Paps & Holland 2018).

Third, in contrast to the relationship between the loss of regulatory genes and the advents of parasitism within the rotifers, flatworms and nematomorphs, no relationship exists between morphological complexity and the number of regulatory protein-encoding genes, whether TFs or RBPs (Fig. 4B; Table 3). It has long been known – and again as we show here (Table 3) – that there is no correlation between an increase to either genome size (Gregory 2001) or to the number of PEGs (Hahn & Wray 2002) and a resulting increase to organismal complexity, the C-value and G-value paradoxes, respectively. Nonetheless, humans do have more *regulatory* genes – both TFS and RBPs (Supp. File 1) – relative to most other animals, and thus the complexity of gene expression could then lead to the organismal complexity like that seen in the vertebrate lineage (e.g., (Carroll 2001; Levine & Tjian 2003; Szathmáry et al. 2001). Because this increase in both TFs and regulatory RBPs is largely due to the two rounds of WGD that occurred early in vertebrate history (Dehal & Boore 2005; Simakov et al. 2020; Lamb 2021; Yu et al. 2024; Marlétaz et al. 2024), and because vertebrates were the only bilaterians to show evidence for WGD events (Ohno 1970), it seemed likely then that WGDs were the ultimate cause of organismal complexity thanks to the “neo-functionalization” of these newly generated paralogues (e.g., (Holland et al. 1994; Escriva et al. 2006; Freeling & Thomas 2006; Putnam et al. 2008; Peer et al. 2009).

However, numerous instances of WGD are now known throughout the animal kingdom that are not accompanied by any change to complexity including the teleost fishes (Amores 1998; Jaillon et al. 2004; Braasch et al. 2016), bdelloid rotifers (Welch et al. 2008; Simion et al. 2021) and horseshoe crabs (Kenny et al. 2016; Nong et al. 2021) analyzed herein (Fig. 1). Importantly, none of these WGDs were suspected and were only discovered when their respective genomes were sequenced, highlighting the disjoint between WGD and correlative morphological complexity. Instead, it now seems clear that at least the initial reason for TF and regulatory RBP retention has more to do with dosage considerations than with neo-functionalization (see (Peterson et al. 2022) and citations therein) and thus are unpredictable given an organism’s relative complexity. In fact, a careful study of *Hox* gene utilization in teleosts in comparison to mice showed that each lineage is characterized by instances of sub- and neo-functionalization that evolved uniquely within each lineage (Yamada et al. 2021) long after the two rounds of WGD. Of course, this model does not preclude subsequent neo-functionalization events (Mantica et al. 2024), but does suggest that the initial reason for genomic retention is much more about retaining the ancestral stoichiometric relationships between the regulator of interest and its regulatory partners and targets rather than unique roles derived quickly after the duplication event(s) to mediate phenotypic changes in morphology (Thompson et al. 2016). Although it remains possible that vertebrates are more complex than other animals that also experienced WGDs like horseshoe crabs thanks to their generation of more regulatory *products* through the complexification of developmental *cis*-regulation, without experimental data showing otherwise, it seems improbable that this would occur in vertebrates despite their having fewer transcriptional and co-transcriptional regulatory *players*. Indeed, when a living fossil like the horseshoe crab whose morphological stasis extends back over 400 million years (Rudkin et al. 2008) possesses more regulatory protein-encoding genes than humans, but yet shares a basic nervous system with other chelicerates and exhibits a relatively simple behavioral repertoire, the role WGDs play in the evolution of complex morphologies – if any – needs to be carefully reconsidered (Kenny et al. 2016).

Which brings us to our fourth and final conclusion: there is a clear difference in the number of post-transcriptional regulatory players (i.e., miRNA families) between morphologically complex organisms like human and octopus versus simpler organisms like horseshoe crabs and bdelloid rotifers (Table 1). In fact, the strongest correlation between any of our individual data sets in the relationship between the number of miRNA families, which are the primary loaders on GPC1, and complexity as measured by estimates of cell types in the adult (Fig. 4A, Table 3), a metric originally proposed by Valentine et al. (1993) and whose general trend is consistent with most recent single-cell RNA sequence analyses (e.g., (Lamanna et al. 2023; Musser et al. 2021; Zeisel et al. 2015; Siebert et al. 2019; Álvarez-Campos et al. 2024; Plass et al. 2018). miRNAs have long been suspected to potentially drive the evolution of novel cell types (Sempere et al. 2006; Lee et al. 2007; Kosik 2010; Berezikov 2011; Zolboot et al. 2021) given the correlative increases to miRNA families at specific points in animal evolution with the cell type that primarily expresses the miRNA (e.g., (Christodoulou et al. 2010; Heimberg et al. 2008). For example, when examining cell-type specific expression of miRNAs in vertebrate-specific blood cells (Patil et al. 2022), the highest expressed cell-type specific miRNA is also a vertebrate-specific miRNA. For example, in neutrophils, the highest expressed cell-type specific miRNA is the vertebrate-specific miRNA Mir-223; in erythrocytes it is the vertebrate-specific pair of co-transcribed miRNAs Mir-451 and Mir-144; in mast cells it is the vertebrate-specific Mir-148; and in B and T lymphocytes it is the vertebrate-specific Mir-150. Even the upstream specification of vertebrate hematopoietic cells uses the vertebrate-specific miRNA Mir-142 (Chen et al. 2004; Merkerova et al. 2008), which remains as the highest expressed miRNA in natural-killer cells. Similarly, in placental epithelial cells, the only cell-type specific miRNAs expressed at or above biologically meaningful levels (i.e., > 1,000 rpm, (Mullokandov et al. 2012) are the therian-specific miRNA Mir-483, and the eutherian-specific miRNAs Mir-127 and Mir-43 (Patil et al. 2022). Thus, at two very different points in evolutionary history, and involving two very different types of tissues, the evolutionary advent of the tissue also corresponds to the evolutionary advent of the cell-type specific miRNA(s).

Importantly, the only example of an obvious increase in morphological complexity within the invertebrates is the coleoid molluscs, the group that includes the octopus and squid. These organisms are famous for their morphologically complex nervous systems (Styfhals et al. 2022) and their remarkable intellectual and behavioral repertoires (Amodio et al. 2019; Gutnick et al. 2023; Schnell et al. 2021). This is also the only known instance of a pulse of miRNA innovation within the non-vertebrates after the divergence of bilaterians from eumetazoans (Supp. Fig. 2), an extensive repertoire of miRNAs that are largely expressed in the adult nervous system (Zolotarov et al. 2022). Other factors often considered causal in cephalopod complexity are singletons not found in other instances of complexification, but often found in other, more simple, organisms. For example, the octopus is characterized by a dramatic increase in the number of C_2_H_2_ zinc finger genes (Albertin et al. 2015), but so too is the morphologically simpler amphioxus that possesses more C_2_H_2_ zinc finger genes than human (Schmitz et al. 2016); coleoids have highly rearranged genomes (Schmidbaur et al. 2022) but so too does the polychaete annelid *D. gyrociliatus* in relation to its near relative *C. teleta* (Martín-Durán et al. 2021). As Carroll (Carroll 2001) emphasized, it is the repeatability of a potential causal factor in different lineages that helps us see meaningful trends, and the only known pulses of miRNA family-level innovation are also the only known instances of increases to morphological complexity in the animal kingdom (Fig. 1); no other correlate is found (Table 3) or known, and no contrary examples have been discovered.

But this raises an interesting question: why does miRNA innovation potentially lead to the evolution of new cell types? Bird (Bird 1995) suggested that the limit to biological complexity might be the efficiency of negative gene regulation as any new cell type requires additional information to reduce the transcriptional noise associated with a new pathway of cellular specification. Although he looked to other potential forms of negative gene regulation, this is exactly what miRNAs achieve during development: they control the accuracy and precision of target gene expression by the unique roles they play in the negative regulation of their target’s mRNA stability and translation (Bartel & Chen 2004; Hornstein & Shomron 2006; Peterson et al. 2009; Wu et al. 2009). These targets are largely the TFs and regulatory RBPs that are involved in organismal development (Zare et al. 2014; Stark et al. 2005), the very same factors considered herein in the construction of genome space (Fig. 1). The presence of these regulatory proteins themselves make no predictions concerning advents of organismal complexity within the Bilateria; only when the activities of this largely conserved repertoire are regulated by taxonomically unique sets of miRNAs do obvious changes to morphology occur. These changes might more readily evolve, at least in part, thanks to the role that decreasing variation plays in increasing heritability (Wu et al. 2009; Peterson et al. 2009). Therefore, although simplification as seen in rotifers and flatworms, and particularly in their parasitic members, is correlated with the loss of all genic types examined including miRNA families (Fig. 1; Table 3), complexity appears to be the result of unusual (and unexplained) pulses of miRNA innovation that allow selection to change protein-expression profiles more effectively, ultimately allowing for the evolution of novel cell types, novel organs, and potentially even novel behaviors.

## Materials and Methods

### Gene Complements

All regulatory protein sequences were initially compiled following Messina et al. (Messina et al. 2004) for human TFs, the curated sequences for fruit fly TFs in Flybase (https://flybase.org/reports/FBgg0000745.htm), Gerstberger et al. (Gerstberger et al. 2014) for human RBPs, and Gamberi et al. (Gamberi et al. 2006) for fruit fly RBPs. Regulatory RBPs were discerned from non-regulatory RBPs by examining the “GO: Biological Function” for each gene and compiling those with clear regulatory roles in either co-transcriptional or post-transcriptional processing in either human or fruit fly (see Supp. File 1, tab 2). Octopus TFs were annotated by searching the proteome with hmmscan program from HMMER v 3.3.2 with --cut_ga option against the list of transcription factor Pfam models (Mistry et al. 2020). To compile the ancestral repertoire of regulatory proteins present in the LCB versus those that evolved afterwards in either the human, fruit fly, or (for the TFs) the octopus lineage, first orthologues of each gene in human were search initially using Ensemble’s “orthologue” function in three other vertebrate species, the mouse *Mus musculus*, the chicken *Gallas gallus* and the spotted gar *Lepisosteus oculatus*, as well as any paralogues identified using Ensemble’s “paralogue” function. Each gene plus all paralogues were then mapped to one of the four gnathostome-specific sub-genomes derived from one of the ∼17 ancestral chordate linkage groups (Lamb 2021; Simakov et al. 2020) (Supp. File 1, tab 1). This allowed us to distinguish between paralogues generated by the two whole-genome duplication (WDG) events in early vertebrate history (Dehal & Boore 2005) from more ancient gene duplications potentially shared with one or more invertebrate taxa.

To search for paralogues in the invertebrate taxa, we used a combination of reciprocal blastp (using the default settings at NCBI) (Tatusov et al. 1997) and phylogenetic analysis (with sequences aligned using Muscle and the phylogeny generated with Neighbor-joining using uncorrected distances using the default settings in Macvector v. 18.2.5) against the publicly available predicted proteins from the sequenced genomes (assembly accession for each species is given in Supp. File 1, tab 5). The predicted proteins for all but three taxa were taken directly from Genbank. The predicted proteins from the acoel *H. miamia* were taken from Ensembl https://metazoa.ensembl.org/Hofstenia_miamia/Info/Index) using the default blastp settings. The predicted proteins from the polyclad flatworm *P. crozieri* (Leite et al. 2022) and the xenoturbellid *X. bocki* (Schiffer et al. 2023) were searched with the query files (Supp. Files 2 and 3) using BLAT. Multiple hits that were not clearly delineated as proteins derived from alternative splicing (e.g., those labeled isoform X1, X2 etc.) were searched against the assembled genome using the default settings in blastn and the total number of loci recorded in Supp. File 1, tab 1. All sequences for all invertebrate taxa minus *D. melanogaster* (which are available at flybase) are available upon request.

The gains and losses of 763 miRNA families found in at least one of the 32 queried species were taken directly from a pre-release version of MirGeneDB v2.2 (Fromm et al. in prep.), which can be viewed here (https://docs.google.com/spreadsheets/d/1LMivoVccN1WVPUFesTVxMWq_E5MCyZDxOmU xu2b4Fww/edit#gid=184772602) and are summarized in Supp. File 1 (tab 3). Finally, the curated 954 BUSCO gene data set (Manni et al. 2021; Simão et al. 2015) was assembled for each of the 32 sampled species (BUSCO v5.4.3, Metazoa node, Supp. File 1, tab 4). Nine of these BUSCO genes overlapped either the TF file or the RBP file (dark red highlighting in tabs 1 and 2) and were removed from the BUSCO file generating a set of 945 non-regulatory and non-redundant housekeeping genes (Supp. File 1, tab 5). Because BUSCO was originally designed to test for the completeness of genomic sequencing, it is possible that any correlations between low BUSCO values and low regulatory values are simply due to the incompleteness of the genomic sequence itself. However, because we used small RNA libraries to curate the miRNA complements for each taxon, genomic completeness can be estimated by documenting the false negatives, i.e., mature miRNA sequences that do not have a corresponding genomic sequence. Of the 2208 miRNA sequences curated for all taxa (minus *D. gyrociliatus* where small RNA reads are not available) we find only 20 false negatives, half of which are miRNAs from the chicken *G. gallus* and are likely found on the incompletely sequenced minichromosomes of the chicken genome. Hence, any correlated absences are likely meaningful and are not due to incomplete genomic sequencing.

### Genome space

The percentage gain and loss of each of the four gene sets for each of the 32 terminal taxa were calculated with respect to the bilaterian last common ancestor (LCB). These values are given in Supp. File 1 (tab 6) and were used for the principal components analysis (PCA). An alternative ancestor was also explored, one where the xenacoelomorphs (*Xenoturbella* and the acoel *H. miamia*) are nested within the deuterostomes and allied with the ambulacrarians (echinoderms and hemichordates). These data are given in Supp. File 1, tab 12 and are also used for a PCA (Supp. File 1, tab 13), but because both analyses result in near-identical results (Supp. Fig. 1) we only show the results with xenacoelomorphs reconstructed as basal bilaterians (see the text for further discussion). PCA was performed on the covariance matrix for the eight genomic variables using Matlab v.2024a (Mathworks Inc.). Variable loadings and percent of total variance explained are given in Table 2. Principal component scores for the first three PCs (i.e., GenomePC1-GenomePC3) were calculated for each taxon and used in subsequent analyses. Pearson product-moment correlations and linear regressions of PCs and original variables were calculated using SAS v.9.4 (SAS Institute).

### Morphospace

The original Deline et al. (Deline et al. 2018) metazoan morphological dataset consists of 212 extant terminal taxa from 34 animal phyla coded with 1,767 discrete characters. The operational taxonomic units (OTUs) varied in rank from family to phylum, but most represented Linnean orders. Deline et al. (Deline et al. 2018) mapped character contingencies to analytically differentiate between missing, absent, and non-applicable characteristics. We culled the full morphological dataset to only include only the OTUs corresponding to the 32 genetically characterized genera in the current study. Overall, the morphological characterization is coarse given the difference in taxonomic rank but with the genomic dataset but captures broad differences in body plans and defining characteristics. In several instances, multiple taxa fell within the same morphological OTU (e.g. Homo and Mus within placental mammals), in those cases both taxa were coded identically in terms of morphology. Even with the course nature of the morphological datasets, the morphology displayed at the generic level was consistent and accurate with that of the higher taxonomic group. The exceptions are *D. gyrociliatus*, which is morphologically distinctive from its associated OTU (Scolecida) and whose coding was based on (Martín-Durán et al., 2021), and the spider mite *T. urticae*, whose coding was altered from the tick *I. scapularis* based on the characters provided in OConnor (OConnor 2009). In total, we added five characters to the original dataset to both capture autapomorphic characters within added taxa (i.e. *D. gyrociliatus*) or to differentiated distinctive taxa within a broad OTU (e.g. ticks and mites) (See Supp File 2, tabs 1 and 2).

The resulting dataset of 32 taxa and 1,769 characters was analyzed following the methods of Deline et al. (2018). The current analysis only differed in methodology from the previous study in the use of an eigenvector-based ordination (PCO) rather than NMDS, to use consistent methods to those utilized with the genomic data. The resulting morphospace is consistent with that of the previous study and the previous study found that morphspaces generated with these different ordination methods were highly correlated (Deline et al., 2018). The morphological analyses were conducted in R (4.3.2) using the cluster (Maechler 2018) and ape (Paradis & Schliep 2018) packages.

### Phylogenetic Independent Contrasts

To generate phylogenetically independent contrasts (PIC) analyses, a data set consisting of 19 concatenated protein sequences was assembled for each of the 32 ingroup taxa as well as the outgroup *Nematostella vectensis*. These proteins were chosen because of their ability to capture the phylogeny (yeast (human (fly, *C.elegans*))) or to not strongly support an alternative arrangement (Mushegian et al. 1998; Wang et al. 1999). A consensus topology was generated (Maclade v. 4.08a, see Supp. File 1, tab 11) and this topology was used by MEGA7 (Kumar et al. 2016) to estimate divergence times. PIC analyses and phylogenetic regressions were performed using the ape, phytools, and nlme packages of R(Revell & Harmon 2022). The use of alternative topologies (e.g., Xenacoelomorpha as ambulacrarian deuterostomes) had no significant effect on the resulting analysis.

## Acknowledgements

G.Z. is supported INPhINIT PhD fellowship from LaCaixa Foundation LCF/BQ/DI21/11860036. PM acknowledges the Spanish “Ministerio de Ciencia, Innovación y Universidades” (GRANT: PID2021-124415NB-I00) for financial support. A. Clark was supported by the Dartmouth James O. Freedman Presidential Scholars program. We thank P. Donoghue, D. Erwin, J. Fyda and B. Runnegar for their comments, T. Lamb for his help with assigning genes to vertebrate sub-genomes, and J. Alkire, J. Courtine, O. Peterson, D. Wang and A. Warren for their careful reading of an earlier version of this manuscript.

## Author Contributions

K.J.P. conceived the project. K.J.P, A.W.C, G.Z., B.D. and B.F. collected and analyzed data. M.A.M. and B.D performed all statistical analyses. K.J.P and P.M wrote the paper with all authors contributing to the interpretation of the results and the final draft of the manuscript.

## Data availability statement

Fasta files for all curated TFs and regulatory RBPs are available from the corresponding author upon request.

## Description of Supplemental Files

**Supplemental Figure 1.** The phylogenetic position of xenacoelomorphs, whether basal bilaterians (left, LCB = last common bilaterian ancestor) or allied with the ambulacrarian deuterostomes (right, LCN = last common nephrozoan ancestor), are very similar (bottom) and has little effect on the resulting genome space analyses. Because of this similarity, and the fact that it requires fewer gene losses if they are basal bilaterians (gold X’s in both analyses), we only show the results with them reconstructed as basal bilaterians.

**Supplemental File 1**. A .xlsx file of all transcription factors (tab 1), RNA binding proteins (tab 2), miRNA families (tab 3) and BUSCO genes (tabs 4-5) for the 32 species considered herein. The percentage of gains and losses of each of these gene categories relative to the bilaterian last common ancestor (LCB) is given in tab 6. The results of the PCA analyses for both genome space and morphospace are given in tabs 7 and 9, respectively. The data for the regression analyses shown in Figure 2 are found in tab 8. The genes used to build the phylogenetic tree for the phylogenetic independent contrasts are listed in tab 10 and the resulting tree and contrasts are given in tab 11. Tabs 12 and 13 explore the affect treating xenacoelomorphs as ambulacrarian deuterostomes and are the data that generated Supp. Fig. 1B.

**Supplemental File 2**. A .txt file (in fasta format) this was used as the query file for transcription factors.

**Supplemental File 3**. A .txt file (in fasta format) this was used as the query file for the regulatory RNA-binding proteins.

**Supplemental File 4**. A .xlsx file of the characters used for the morphospace analyses for the 32 taxa considered herein. Tab 1 gives the coding for each of the 1767 characters whose descriptions and character states are found in Tab 2 including the coding for *D. gyrociliatus*, which was not included as one of the terminal taxa in the original Deline et al. study. Note that the codings for mouse and human, as well as zebrafish and *Tetraodon*, are identical given the taxonomic levels analyzed in the Deline et al. (2018) study.

**Supplemental File 5**. The alignment of the 19 concatenated protein sequences as a .txt file (in nexus format) for the 32 considered ingroup species as well as the outgroup *N. vectensis* used for the phylogenetic independent contrasts analyses (Supp. File 1, tab 11). The list of the 19 proteins as well as their representation across the ingroup species are given in Supp. File 1, tab 10.

## References

Albalat R, Cañestro C. 2016. Evolution by gene loss. Nat. Rev. Genet. 17:379–391. doi: 10.1038/nrg.2016.39.

Albertin CB et al. 2015. The octopus genome and the evolution of cephalopod neural and morphological novelties. Nature. 524:220–224. doi: 10.1038/nature14668.

Albertin CB, Katz PS. 2023. Evolution of cephalopod nervous systems. Curr. Biol. 33:R1087– R1091. doi: 10.1016/j.cub.2023.08.092.

Álvarez-Campos P et al. 2024. Annelid adult cell type diversity and their pluripotent cellular origins. Nat. Commun. 15:3194. doi: 10.1038/s41467-024-47401-6.

Amodio P et al. 2019. Grow Smart and Die Young: Why Did Cephalopods Evolve Intelligence? Trends Ecol. Evol. 34:45–56. doi: 10.1016/j.tree.2018.10.010.

Amores A. 1998. Zebrafish hox clusters and vertebrate genome evolution. Science. 282:1711– 1714. doi: 10.1126/science.282.5394.1711.

Anantharaman V, Koonin EV, Aravind L. 2002. Comparative genomics and evolution of proteins involved in RNA metabolism. Nucleic Acids Res. 30:1427–1464. doi: 10.1093/nar/30.7.1427.

Bartel DP, Chen C-Z. 2004. Micromanagers of gene expression: the potentially widespread influence of metazoan microRNAs. Nat Rev Genet. 5:396–400. doi: 10.1038/nrg1328.

Bell G, Mooers AO. 1997. Size and complexity among multicellular organisms. Biol. J. Linn. Soc. 60:345–363. doi: 10.1111/j.1095-8312.1997.tb01500.x.

Berezikov E. 2011. Evolution of microRNA diversity and regulation in animals. Nat Rev Genet. 12:846–860. doi: 10.1038/nrg3079.

Berriman M et al. 2009. The genome of the blood fluke Schistosoma mansoni. Nature. 460:352–358. doi: 10.1038/nature08160.

Bird AP. 1995. Gene number, noise-reduction and biological complexity. Trends. Genet. 11:94–100. doi: 10.1016/s0168-9525(00)89009-5.

Bonner JT. 2004. Perspective: the size-complexity rule. Evolution. 58:1883–1890. doi: 10.1111/j.0014-3820.2004.tb00476.x.

Braasch I et al. 2016. The spotted gar genome illuminates vertebrate evolution and facilitates human-teleost comparisons. Nat. Genet. 48:427–437. doi: 10.1038/ng.3526.

Brabec J, Salomaki ED, Kolísko M, Scholz T, Kuchta R. 2023. The evolution of endoparasitism and complex life cycles in parasitic platyhelminths. Curr. Biol. doi: 10.1016/j.cub.2023.08.064.

Cannon JT et al. 2016. Xenacoelomorpha is the sister group to Nephrozoa. Nature. 530:89–93. doi: 10.1038/nature16520.

Carroll SB. 2001. Chance and necessity: the evolution of morphological complexity and diversity. Nature. 409:1102–1109. doi: 10.1038/35059227.

Carroll SB. 2008. Evo-devo and an expanding evolutionary synthesis: a genetic theory of morphological evolution. Cell. 134:25–36. doi: 10.1016/j.cell.2008.06.030.

Chen C-Z, Li L, Lodish HF, Bartel DP. 2004. MicroRNAs modulate hematopoietic lineage differentiation. Science. 303:83–86. doi: 10.1126/science.1091903.

Christodoulou F et al. 2010. Ancient animal microRNAs and the evolution of tissue identity. Nature. 1–7. doi: 10.1038/nature08744.

Coghlan A et al. 2019. Comparative genomics of the major parasitic worms. Nat. Genet. 51:163–174. doi: 10.1038/s41588-018-0262-1.

Consortium SUGS et al. 2006. The genome of the sea urchin Strongylocentrotus purpuratus. Science. 314:941–952. doi: 10.1126/science.1133609.

Cunha TJ, Medeiros BAS de, Lord A, Sørensen MV, Giribet G. 2023. Rampant loss of universal metazoan genes revealed by a chromosome-level genome assembly of the parasitic Nematomorpha. Curr. Biol. 33:3514–3521.e4. doi: 10.1016/j.cub.2023.07.003.

De S et al. 2023. A high-quality Ixodes scapularis genome advances tick science. Nat. Genet. 55:301–311. doi: 10.1038/s41588-022-01275-w.

Degnan BM, Vervoort M, Larroux C, Richards GS. 2009. Early evolution of metazoan transcription factors. Curr Opin Genet Dev. 19:591–599. doi: 10.1016/j.gde.2009.09.008.

Dehal P, Boore J. 2005. Two rounds of whole genome duplication in the ancestral vertebrate. Plos Biol. 3:1700–1708. doi: 10.1371/journal.pbio.0030314.

Deline B et al. 2018. Evolution of metazoan morphological disparity. Proc. Natl. Acad. Sci. U.S.A. 29:201810575–10. doi: 10.1073/pnas.1810575115.

Devor EJ, Peek AS. 2008. miRNA Profile of a Triassic Common Mammalian Ancestor and PremiRNA Evolution in the Three Mammalian Reproductive Lineages. TOGENJ. 1:22–32. doi: 10.2174/1875693x00801010022.

Duruz J et al. 2020. Acoel Single-Cell Transcriptomics: Cell Type Analysis of a Deep Branching Bilaterian. Mol. Biol. Evol. 38:1888–1904. doi: 10.1093/molbev/msaa333.

Erwin DH. 2007. Disparity: Morphological pattern and developmental context. Palaeontology. 50:57–73. doi: 10.1111/j.1475-4983.2006.00614.x.

Escriva H et al. 2006. Neofunctionalization in vertebrates: the example of retinoic acid receptors. PLoS Genet. 2:e102. doi: 10.1371/journal.pgen.0020102.

Felsenstein J. 1985. Phylogenies and the Comparative Method. Am. Nat. 125:1–15. doi: 10.1086/284325.

Fernández R, Gabaldon T. 2020. Gene gain and loss across the metazoan tree of life. Nat. Ecol. Evol. 1–12. doi: 10.1038/s41559-019-1069-x.

Foote M. 1997. The evolution of morphological diversity. Annu. Rev. Ecol. Syst. 28:129–152. doi: 10.1146/annurev.ecolsys.28.1.129.

Freeling M, Thomas BC. 2006. Gene-balanced duplications, like tetraploidy, provide predictable drive to increase morphological complexity. Genome Res. 16:805–814. doi: 10.1101/gr.3681406.

Fromm B. 2024. A renaissance of microRNAs as taxonomic and phylogenetic markers in animals. Zoöl. Scr. doi: 10.1111/zsc.12684.

Fromm B et al. 2022. MirGeneDB 2.1: toward a complete sampling of all major animal phyla. Nucleic Acids Res. 50:D204–D210. doi: 10.1093/nar/gkab1101.

Gamberi C, Johnstone O, Lasko P. 2006. Drosophila RNA Binding Proteins. Int. Rev. Cytol. 248:43–139. doi: 10.1016/s0074-7696(06)48002-5.

Gehrke AR et al. 2019. Acoel genome reveals the regulatory landscape of whole-body regeneration. Science. 363. doi: 10.1126/science.aau6173.

Gerstberger S, Hafner M, Tuschl T. 2014. A census of human RNA-binding proteins. Nat Rev Genet. 15:829–845. doi: 10.1038/nrg3813.

Grbić M et al. 2011. The genome of Tetranychus urticae reveals herbivorous pest adaptations. Nature. 479:487–492. doi: 10.1038/nature10640.

Gregory TR. 2001. Coincidence, coevolution, or causation? DNA content, cell size, and the C-value enigma. Biol. Rev. Camb. Philos. Soc. 76:65–101. doi: 10.1017/s1464793100005595.

Griffiths JS et al. 2022. A draft reference genome of the red abalone, Haliotis rufescens, for conservation genomics. J. Hered. 113:673–680. doi: 10.1093/jhered/esac047.

Guijarro-Clarke C, Holland PWH, Paps J. 2020. Widespread patterns of gene loss in the evolution of the animal kingdom. Nat. Ecol. Evol. 1–7. doi: 10.1038/s41559-020-1129-2.

Gutnick T, Rokhsar DS, Kuba MJ. 2023. Cephalopod behaviour. Curr. Biol. 33:R1083–R1086. doi: 10.1016/j.cub.2023.08.094.

Hagemann L, Mauer KM, Hankeln T, Schmidt H, Herlyn H. 2023. Nuclear genome annotation of wheel animals and thorny-headed worms: inferences about the last common ancestor of Syndermata (Rotifera s.l.). Hydrobiologia. 1–18. doi: 10.1007/s10750-023-05268-6.

Hahn M, Wray G. 2002. The g-value paradox. Evol Dev. 4:73–75.

Heimberg AM, Sempere LF, Moy VN, Donoghue PCJ, Peterson KJ. 2008. MicroRNAs and the advent of vertebrate morphological complexity. Proc. Natl. Acad. Sci. USA. 105:2946–2950. doi: 10.1073/pnas.0712259105.

Herlyn H et al. 2024. Substantial hierarchical reductions of genetic and morphological traits in the evolution of rotiferan parasites. bioRxiv. 2024.08.01.605096. doi: 10.1101/2024.08.01.605096.

Hill MS, Zande PV, Wittkopp PJ. 2021. Molecular and evolutionary processes generating variation in gene expression. Nat. Rev. Genet. 22:203–215. doi: 10.1038/s41576-020-00304-w.

Holland PW, Garcia-Fernandez J, Williams NA, Sidow A. 1994. Gene duplications and the origins of vertebrate development. Dev Suppl. 125–133. http://eutils.ncbi.nlm.nih.gov/entrez/eutils/elink.fcgi?dbfrom=pubmed&id=7579513&retmode=ref&cmd=prlinks.

Holt RA et al. 2002. The Genome Sequence of the Malaria Mosquito Anopheles gambiae. Science. 298:129–149. doi: 10.1126/science.1076181.

Hornstein E, Shomron N. 2006. Canalization of development by microRNAs. Nat. Genet. 38:S20–S24. doi: 10.1038/ng1803.

Hulett RE et al. 2023. Acoel single-cell atlas reveals expression dynamics and heterogeneity of adult pluripotent stem cells. Nat. Commun. 14:2612. doi: 10.1038/s41467-023-38016-4.

Jackson AP. 2015. The evolution of parasite genomes and the origins of parasitism. Parasitology. 142:S1–S5. doi: 10.1017/s0031182014001516.

Jaillon O et al. 2004. Genome duplication in the teleost fish Tetraodon nigroviridis reveals the early vertebrate proto-karyotype. Nature. 431:946–957. doi: 10.1038/nature03025.

Jiang Y, Xu C. 2010. The calculation of information and organismal complexity. Biol. Direct. 5:59. doi: 10.1186/1745-6150-5-59.

Juravel K, Porras L, Höhna S, Pisani D, Wörheide G. 2023. Exploring genome gene content and morphological analysis to test recalcitrant nodes in the animal phylogeny. PLOS ONE. 18:e0282444. doi: 10.1371/journal.pone.0282444.

Kenny NJ et al. 2016. Ancestral whole-genome duplication in the marine chelicerate horseshoe crabs. Heredity. 116:190–199. doi: 10.1038/hdy.2015.89.

Kerner P, Degnan SM, Marchand L, Degnan BM, Vervoort M. 2011. Evolution of RNA-Binding Proteins in Animals: Insights from Genome-Wide Analysis in the Sponge Amphimedon queenslandica. Mol. Biol. Evol. 28:2289–2303. doi: 10.1093/molbev/msr046.

Kosik KS. 2010. MicroRNAs and Cellular Phenotypy. Cell. 143:21–26. doi: 10.1016/j.cell.2010.09.008.

Kumar S, Stecher G, Tamura K. 2016. MEGA7: Molecular Evolutionary Genetics Analysis Version 7.0 for Bigger Datasets. Mol. Biol. Evol. 33:1870–1874. doi: 10.1093/molbev/msw054.

Lamanna F et al. 2023. A lamprey neural cell type atlas illuminates the origins of the vertebrate brain. Nat. Ecol. Evol. 7:1714–1728. doi: 10.1038/s41559-023-02170-1.

Lamb TD. 2021. Analysis of paralogons, origin of the vertebrate karyotype, and ancient chromosomes retained in extant species. Genome Biol Evol. 13. doi: 10.1093/gbe/evab044.

Lambert SA et al. 2018. The human transcription factors. Cell. 172:650–665. doi: 10.1016/j.cell.2018.01.029.

Langschied F, Leisegang MS, Brandes RP, Ebersberger I. 2023. ncOrtho: efficient and reliable identification of miRNA orthologs. Nucleic Acids Res. doi: 10.1093/nar/gkad467.

Lee C-T, Risom T, Strauss WM. 2007. Evolutionary Conservation of MicroRNA Regulatory Circuits: An Examination of MicroRNA Gene Complexity and Conserved MicroRNA-Target Interactions through Metazoan Phylogeny. DNA and Cell Biology. 26:209–218. doi: 10.1089/dna.2006.0545.

Leite DJ, Piovani L, Telford MJ. 2022. Genome assembly of the polyclad flatworm Prostheceraeus crozieri. Genome Biol. Evol. 14:evac133. doi: 10.1093/gbe/evac133.

Levine M, Tjian R. 2003. Transcription regulation and animal diversity. Nature. 424:147–151. doi: 10.1038/nature01763.

Lewontin RC. 1974. *The Genetic Basis of Evolutionary Change*. Columbia University Press: New York City.

Luo Y-J et al. 2015. The Lingula genome provides insights into brachiopod evolution and the origin of phosphate biomineralization. Nat Commun. 6:8301. doi: 10.1038/ncomms9301.

Maechler M. 2018. Cluster: cluster analysis basics and extensions.

Manni M, Berkeley MR, Seppey M, Simão FA, Zdobnov EM. 2021. BUSCO update: novel and streamlined workflows along with broader and deeper phylogenetic coverage for scoring of eukaryotic, prokaryotic, and viral genomes. Mol. Biol. Evol. 38:msab199-. doi: 10.1093/molbev/msab199.

Mantica F et al. 2024. Evolution of tissue-specific expression of ancestral genes across vertebrates and insects. Nat. Ecol. Evol. 1–14. doi: 10.1038/s41559-024-02398-5.

Marlétaz F et al. 2024. The hagfish genome and the evolution of vertebrates. Nature. 627:811– 820. doi: 10.1038/s41586-024-07070-3.

Martín-Durán JM et al. 2021. Conservative route to genome compaction in a miniature annelid. Nat. Ecol. Evol. 5:231–242. doi: 10.1038/s41559-020-01327-6.

Mauer K et al. 2020. The genome, transcriptome, and proteome of the fish parasite Pomphorhynchus laevis (Acanthocephala). PLoS ONE. 15:e0232973. doi: 10.1371/journal.pone.0232973.

Mauer KM et al. 2021. Genomics and transcriptomics of epizoic Seisonidea (Rotifera, syn. Syndermata) reveal strain formation and gradual gene loss with growing ties to the host. BMC Genom. 22:604. doi: 10.1186/s12864-021-07857-y.

McShea DW. 2000. Functional complexity in organisms: Parts as proxies. Biol. Philos. 15:641– 668. doi: 10.1023/a:1006695908715.

McShea DW. 1996. Metazoan complexity and evolution: Is there a trend? Evolution. 50:477–492. doi: 10.1111/j.1558-5646.1996.tb03861.x.

McShea DW, Brandon Robert N. 2010. *Biology’s First Law: The Tendency for Diversity and Complexity to Increase in Evolutionary Systems*. University of Chicago Press: Chicago.

Merkerova M, Belickova M, Bruchova H. 2008. Differential expression of microRNAs in hematopoietic cell lineages. Eur. J. Haematol. 81:304–310. doi: 10.1111/j.1600-0609.2008.01111.x.

Messina DN, Glasscock J, Gish W, Lovett M. 2004. An ORFeome-based Analysis of Human Transcription Factor Genes and the Construction of a Microarray to Interrogate Their Expression. Genome Res. 14:2041–2047. doi: 10.1101/gr.2584104.

Mistry J et al. 2020. Pfam: The protein families database in 2021. Nucleic Acids Res. 49:D412– D419. doi: 10.1093/nar/gkaa913.

Mulhair PO, McCarthy CGP, Siu-Ting K, Creevey CJ, O’Connell MJ. 2022. Filtering artifactual signal increases support for Xenacoelomorpha and Ambulacraria sister relationship in the animal tree of life. Curr Biol. doi: 10.1016/j.cub.2022.10.036.

Mullokandov G et al. 2012. High-throughput assessment of microRNA activity and function using microRNA sensor and decoy libraries. Nat. Methods. 9:840–846. doi: 10.1038/nmeth.2078.

Mushegian AR, Garey JR, Martin J, Liu LX. 1998. Large-scale taxonomic profiling of eukaryotic model organisms: A comparison of orthologous proteins encoded by the human, fly, nematode, and yeast genomes. Genome Res. 8:590–598. doi: 10.1101/gr.8.6.590.

Musser JM et al. 2021. Profiling cellular diversity in sponges informs animal cell type and nervous system evolution. Science. 374:717–723. doi: 10.1126/science.abj2949.

Neafsey DE et al. 2015. Highly evolvable malaria vectors: The genomes of 16 Anopheles mosquitoes. Science. 347:1258522. doi: 10.1126/science.1258522.

Nong W et al. 2021. Horseshoe crab genomes reveal the evolution of genes and microRNAs after three rounds of whole genome duplication. Communications Biology. 1–11. doi: 10.1038/s42003-020-01637-2.

OConnor BM. 2009. Mites. In: Encyclopedia of Insects. pp. 643–649.

Ohno S. 1970. *Evolution by Gene Duplication*. Berlin, New York, Springer-Verlag.

Paps J, Holland PWH. 2018. Reconstruction of the ancestral metazoan genome reveals an increase in genomic novelty. Nat Commun. 1–8. doi: 10.1038/s41467-018-04136-5.

Paradis E, Schliep K. 2018. ape 5.0: an environment for modern phylogenetics and evolutionary analyses in R. Bioinformatics. 35:526–528. doi: 10.1093/bioinformatics/bty633.

Patil AH, Baran A, Brehm ZP, McCall MN, Halushka MK. 2022. A curated human cellular microRNAome based on 196 primary cell types. GigaScience. 11:giac083. doi: 10.1093/gigascience/giac083.

Peer YV de, Maere S, Meyer A. 2009. The evolutionary significance of ancient genome duplications. Nature. 1–8. doi: 10.1038/nrg2600.

Peter IS, Davidson EH. 2015. *Genomic Control Process: Development and Evolution*. Academic Press: Amsterdam.

Peterson KJ et al. 2022. MicroRNAs as indicators into the causes and consequences of whole-genome duplication events. Mol. Biol. Evol. 39:msab344. doi: 10.1093/molbev/msab344.

Peterson KJ, Dietrich MR, McPeek MA. 2009. MicroRNAs and metazoan macroevolution: insights into canalization, complexity, and the Cambrian explosion. Bioessays. 31:736–747. doi: 10.1002/bies.200900033.

Philippe H et al. 2011. Acoelomorph flatworms are deuterostomes related to Xenoturbella. Nature. 470:255–258. doi: 10.1038/nature09676.

Plass M et al. 2018. Cell type atlas and lineage tree of a whole complex animal by single-cell transcriptomics. Science. 360. doi: 10.1126/science.aaq1723.

Poulin R. 2011. Chapter 1 The Many Roads to Parasitism A Tale of Convergence. Adv. Parasitol. 74:1–40. doi: 10.1016/b978-0-12-385897-9.00001-x.

Poulin R, Randhawa HS. 2015. Evolution of parasitism along convergent lines: from ecology to genomics. Parasitology. 142:S6–S15. doi: 10.1017/s0031182013001674.

Putnam NH et al. 2008. The amphioxus genome and the evolution of the chordate karyotype. Nature. 453:1064–U3. doi: 10.1038/nature06967.

Revell LJ, Harmon LJ. 2022. *Phylogenetic comparative methods in R*. Princeton University Press: Princeton.

Richter DJ, Fozouni P, Eisen MB, King N. 2018. Gene family innovation, conservation and loss on the animal stem lineage. eLife. 7:e34226. doi: 10.7554/elife.34226.

Robertson HE et al. 2024. Single cell atlas of Xenoturbella bocki highlights limited cell-type complexity. Nat. Commun. 15:2469. doi: 10.1038/s41467-024-45956-y.

Rouse GW, Wilson NG, Carvajal JI, Vrijenhoek RC. 2016. New deep-sea species of Xenoturbella and the position of Xenacoelomorpha. Nature. 530:94–97. doi: 10.1038/nature16545.

Rudkin DM, Young GA, Nowlan GS. 2008. The oldest horseshoe crab: a new xiphosurid from late Ordovician Konservat-Lagerstätten deposits, Manitoba, Canada. Palaeontology. 51:1–9. doi: 10.1111/j.1475-4983.2007.00746.x.

Runnegar B. 1987. Rates and modes of evolution in the Mollusca. In: Rates of Evolution. Campell, KSW & Day, MF, editors. Routledge: London pp. 39–60.

Schiffer PH et al. 2023.The slowly evolving genome of the xenacoelomorph worm Xenoturbella bocki. bioRxiv. 2022.06.24.497508. doi: 10.1101/2022.06.24.497508.

Schmidbaur H et al. 2022. Emergence of novel cephalopod gene regulation and expression through large-scale genome reorganization. Nat Commun. 13:2172–11. doi: 10.1038/s41467-022-29694-7.

Schmitz JF, Zimmer F, Bornberg-Bauer E. 2016. Mechanisms of transcription factor evolution in Metazoa. Nucleic Acids Res. 44:6287–6297. doi: 10.1093/nar/gkw492.

Schnell AK, Amodio P, Boeckle M, Clayton NS. 2021. How intelligent is a cephalopod? Lessons from comparative cognition. Biol. Rev. 96:162–178. doi: 10.1111/brv.12651.

Sempere LF, Cole CN, McPeek MA, Peterson KJ. 2006. The phylogenetic distribution of metazoan microRNAs: Insights into evolutionary complexity and constraint. J. Exp. Zool. 306B:575–588. doi: 10.1002/jez.b.21118.

Siebert S et al. 2019. Stem cell differentiation trajectories in Hydra resolved at single-cell resolution. Science. 365. doi: 10.1126/science.aav9314.

Simakov O et al. 2020. Deeply conserved synteny resolves early events in vertebrate evolution. Nat. Ecol. Evol. 1–22. doi: 10.1038/s41559-020-1156-z.

Simakov O et al. 2015. Hemichordate genomes and deuterostome origins. Nature. 527:459–465. doi: 10.1038/nature16150.

Simakov O et al. 2013. Insights into bilaterian evolution from three spiralian genomes. Nature. 493:526–531. doi: 10.1038/nature11696.

Simão FA, Waterhouse RM, Ioannidis P, Kriventseva EV, Zdobnov EM. 2015. BUSCO: assessing genome assembly and annotation completeness with single-copy orthologs. Bioinformatics. 31:3210–3212. doi: 10.1093/bioinformatics/btv351.

Simion P et al. 2021. Chromosome-level genome assembly reveals homologous chromosomes and recombination in asexual rotifer Adineta vaga. Sci. Adv. 7:eabg4216. doi: 10.1126/sciadv.abg4216.

Srivastava M et al. 2010. The Amphimedon queenslandica genome and the evolution of animal complexity. Nature. 1–8. doi: 10.1038/nature09201.

Stark A, Brennecke J, Bushati N, Russell RB, Cohen SM. 2005. Animal microRNAs confer robustness to gene expression and have a significant impact on 3′UTR evolution. Cell. 123:1133–1146. doi: 10.1016/j.cell.2005.11.023.

Styfhals R et al. 2022. Cell type diversity in a developing octopus brain. Nat Commun. 13:7392. doi: 10.1038/s41467-022-35198-1.

Szathmáry E, Jordán F, Pál C. 2001. Can Genes Explain Biological Complexity? Science. 292:1315–1316. doi: 10.1126/science.1060852.

Takatori N et al. 2008. Comprehensive survey and classification of homeobox genes in the genome of amphioxus, Branchiostoma floridae. Dev Genes Evol. 218:579–590. doi: 10.1007/s00427-008-0245-9.

Tarver JE et al. 2013. miRNAs: small genes with big potential in metazoan phylogenetics. Mol. Biol. Evol. 30:2369–2382. doi: 10.1093/molbev/mst133.

Tarver JE et al. 2018. Well-annotated microRNAomes do not evidence pervasive miRNA loss. Genome Biol Evol. 10:1457–1470. doi: 10.1093/gbe/evy096.

Tatusov RL, Koonin EV, Lipman DJ. 1997. A Genomic Perspective on Protein Families. Science. 278:631–637. doi: 10.1126/science.278.5338.631.

Thompson A, Zakon HH, Kirkpatrick M. 2016. Compensatory drift and the evolutionary dynamics of dosage-sensitive duplicate genes. Genetics. 202:765–774. doi: 10.1534/genetics.115.178137.

Treiber T et al. 2017. A compendium of RNA-binding proteins that regulate microRNA biogenesis. Mol. Cell. 66:270–284.e13. doi: 10.1016/j.molcel.2017.03.014.

Tsai IJ et al. 2013. The genomes of four tapeworm species reveal adaptations to parasitism. Nature. 496:57–63. doi: 10.1038/nature12031.

Tupler R, Perini G, Green MR. 2001. Expressing the human genome. Nature. 409:832–833. doi: 10.1038/35057011.

Valentine JW, Collins AG, Meyer CP. 1994. Morphological complexity increase in metazoans. Paleobiology. 20:131–142. doi: 10.1017/s0094837300012641.

Vogel C, Chothia C. 2006. Protein family expansions and biological complexity. PLoS Comp Biol. 2:370–382. doi: 10.1371/journal.pcbi.0020048.

Wang DY, Kumar S, Hedges SB. 1999. Divergence time estimates for the early history of animal phyla and the origin of plants, animals and fungi. Proc. Biol. Sci. 266:163–171. doi: 10.1098/rspb.1999.0617.

Weinstein SB, Kuris AM. 2016. Independent origins of parasitism in Animalia. Biol. Lett. 12:20160324. doi: 10.1098/rsbl.2016.0324.

Welch DBM, Welch JLM, Meselson M. 2008. Evidence for degenerate tetraploidy in bdelloid rotifers. Proc. Natl. Acad. Sci. U.S.A. 105:5145–5149. doi: 10.1073/pnas.0800972105.

Wolf YI, Koonin EV. 2013. Genome reduction as the dominant mode of evolution. BioEssays. 35:829–837. doi: 10.1002/bies.201300037.

Wray GA. 2007. The evolutionary significance of cis-regulatory mutations. Nat Rev Genet. 8:206–216. doi: 10.1038/nrg2063.

Wu C-I, Shen Y, Tang T. 2009. Evolution under canalization and the dual roles of microRNAs: a hypothesis. Genome Res. 19:734–743. doi: 10.1101/gr.084640.108.

Yamada K et al. 2021. An atlas of seven zebrafish hox cluster mutants provides insights into sub/neofunctionalization of vertebrate Hox clusters. Development. 148. doi: 10.1242/dev.198325.

Ye Z et al. 2017. A New Reference Genome Assembly for the Microcrustacean Daphnia pulex. G3: GenesGenomesGenet. 7:1405–1416. doi: 10.1534/g3.116.038638.

Young JZ. 1963. The number and sizes of nerve cells in Octopus. Proc. Zoöl. Soc. Lond. 140:229–254. doi: 10.1111/j.1469-7998.1963.tb01862.x.

Yu D et al. 2024. Hagfish genome elucidates vertebrate whole-genome duplication events and their evolutionary consequences. Nat. Ecol. Evol. 1–17. doi: 10.1038/s41559-023-02299-z.

Zare H, Khodursky A, Sartorelli V. 2014. An evolutionarily biased distribution of miRNA sites toward regulatory genes with high promoter-driven intrinsic transcriptional noise. Bmc Evol Biol. 14:74. doi: 10.1186/1471-2148-14-74.

Zeisel A et al. 2015. Cell types in the mouse cortex and hippocampus revealed by single-cell RNA-seq. Science. 347:1138–1142. doi: 10.1126/science.aaa1934.

Zheng H et al. 2013. The genome of the hydatid tapeworm Echinococcus granulosus. Nat. Genet. 45:1168–1175. doi: 10.1038/ng.2757.

Zolboot N, Du JX, Zampa F, Lippi G. 2021. MicroRNAs Instruct and Maintain Cell Type Diversity in the Nervous System. Front. Mol. Neurosci. 14:646072. doi: 10.3389/fnmol.2021.646072.

Zolotarov G et al. 2022. MicroRNAs are deeply linked to the emergence of the complex octopus brain. Sci. Adv. 8:eadd9938. doi: 10.1126/sciadv.add9938.

