## Supplementary figures and images for "Capturing changes to animal complexity from quantifiable patterns in genomic data"

### Supplemental Figure 1

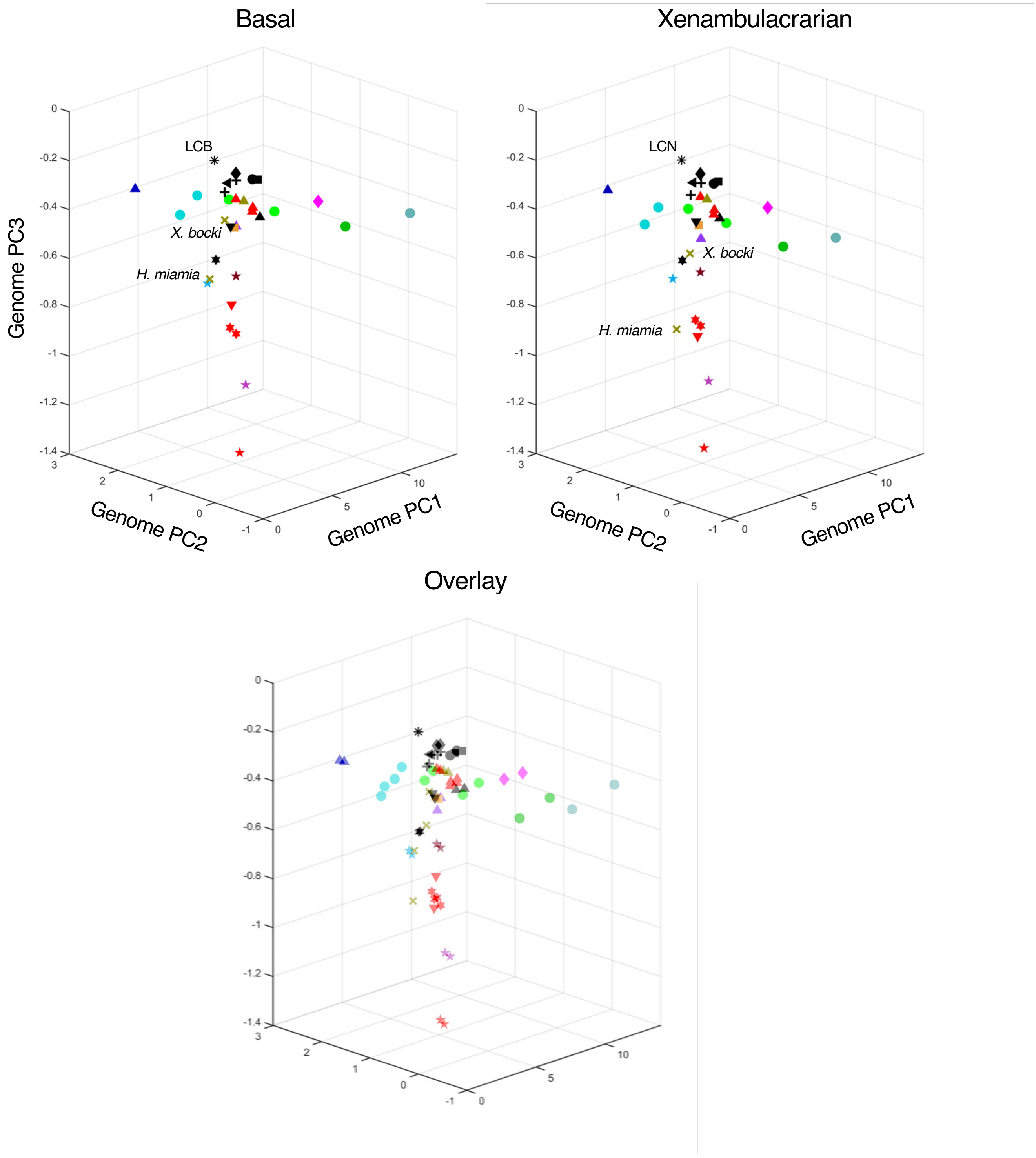
